# HALO: Hierarchical Causal Modeling for Single Cell Multi-Omics Data

**DOI:** 10.1101/2022.10.17.512602

**Authors:** Haiyi Mao, Minxue Jia, Marissa Di, Eleanor Valenzi, Xiaoyu Tracy Cai, Robert Lafyatis, Kun Zhang, Panayiotis V. Benos

## Abstract

Though open chromatin may promote active transcription, gene expression responses may not be directly coordinated with changes in chromatin accessibility. Most existing methods for single-cell multi-omics data focus only on learning stationary and shared information among these modalities, overlooking modality-specific information delineating cellular states and dynamics resulting from causal relations among modalities. To account for this, the epigenome and transcriptome relationship can be characterized in relation to time as “coupled” (changing dependently) or “decoupled” (changing independently). We propose the framework HALO, which adopts a causal approach to model these temporal causal relations on two levels. On the representation level, HALO factorizes these two modalities into both coupled and decoupled latent representations, identifying the dynamic interplay between chromatin accessibility and transcription through temporal modulations in the latent space. On the individual gene level, HALO matches gene-peak pairs and characterizes changing dynamics between gene expression and local peaks with time. HALO reveals bipotency in a subset of AT2 cells that exhibit different decisions in lineage specification between systemic sclerosis (SSc) and normal conditions. We demonstrate that using coupled and decoupled information, HALO discovers analogous biological functions between modalities, distinguishes epigenetic factors for lineage specification, and identifies temporal *cis*-regulation interactions relevant to cellular differentiation and complex human diseases.

## Introduction

Single-cell multi-omics technologies have revolutionized our understanding of cellular heterogeneity and complexity, enabling the simultaneous measurement of diverse molecular layers such as transcriptomics, epigenomics, and proteomics within individual cells. These technologies provide a unique opportunity to dissect the regulatory relationships underlying cellular functions and phenotypes. However, integration and analysis of such heterogeneous data types remain challenging, for example in distinguishing the effects of chromatin accessibility on expression across varying cellular states and over time. To address these challenges, we developed HALO (**H**ierarchical C**A**usal Modeling for Single Cell Mu**L**ti-**O**mics Data), a computational framework designed to model the causal relationships within single-cell co-profiled multi-omic data. The key hypothesis of HALO is that changes in chromatin accessibility causally influence gene expression. By leveraging measurements of transcriptomics and chromatin accessibility data from single-cell RNA sequencing (scRNA-seq) and single-cell ATAC sequencing (scATAC-seq), HALO enables a comprehensive analysis of the causal interactions that dictate cell state, function, and regulatory mechanisms.

Chromatin structure plays a crucial role in regulating the ability of transcriptional machinery to access DNA and activate gene expression. Open chromatin facilitates the binding of transcription factors, the recruitment of RNA polymerases, and the initiation of transcription. Consequently, gene expression and chromatin accessibility often correlate and exhibit similar dynamics across different cellular states or over time. However, they do not always display the same patterns due to several biological regulatory factors. For example, chromatin can be accessible, but the associated gene or regulatory region may not be immediately transcribed—a state known as chromatin priming. This primed state allows cells to respond more quickly to environmental changes or developmental signals. In this state, cells have the potential to differentiate into various cell types and adapt to different conditions^1^. Additionally, gene expression can be controlled post-transcriptionally through mechanisms such as mRNA stability, degradation, or translation efficiency. Meanwhile, active chromatin remodeling may occur independently of transcriptional changes, preparing the chromatin landscape for future gene activation or repression^2^. Previous representation learning methods for single-cell multi-omics data implicitly rely on the strict biological assumption that chromatin accessibility and active transcription are synchronized^3–6^, though some recent methods consider the case where chromatin opening precedes transcription initiation^2,7^. In contrast to these works, HALO decomposes the relations between chromatin accessibility and gene expression into coupled and decoupled cases. In the coupled case, chromatin accessibility and gene expression exhibit correlated changes over time, while in the decoupled case, they change independently over time. Due to the high dimensionality and sparsity of single cell genomics data, HALO operates on two levels of analysis: the low-dimensional latent representation level and the individual gene level, incorporating real time points/estimated latent time to account for temporal dynamics in cellular development. The framework is equipped with an interpretable neural network to provide biological meaning to the latent representations. Moreover, HALO employs Granger causality to assess context-specific distal cis-regulation in cases where, despite associated chromatin regions becoming more accessible, gene transcription does not increase correspondingly^7^. We observe these situations frequently occur when the chromatin regions overlap with super enhancer regions.

## Results

### HALO: a causal machine learning framework to model the interactions between chromatin accessibility and gene expression

HALO models the co-assayed gene expression and chromatin accessibility in a low-dimensional latent space as well as individual gene and its linked peaks (open chromatin regions) level through a causal lens. In our study, we establish a framework to analyze the causal relationships between jointly profiled scRNA-seq and single-cell ATAC sequencing (scATAC-seq) data, incorporating temporal information. We hypothesize that scATAC-seq data, which is indicative of chromatin accessibility, causally precedes scRNA-seq data due to open chromatin regions influencing gene transcription. Specifically, we distinguish between two types of causal interactions: 1) In the **coupled** case, gene expression and chromatin accessibility exhibit *dependent* changes over time, indicating they are influenced by shared unknown confounders. 2) In the **decoupled** case, certain gene expressions and their local peak patterns change *independently* over time, suggesting the presence of distinct, unknown causal factors affecting gene expressions. This approach extends the concept of time-lagging observed in previous studies^2,8^ to a broader framework, encompassing mechanisms that change independently or dependently over time; for example, chromatin open regions may become more accessible while gene expression level remains stable. This approach aims to elucidate causal relationships between gene expression and chromatin accessibility at both the levels of representations and individual genes, as depicted in **Figure 1A**.

**Figure 1.**
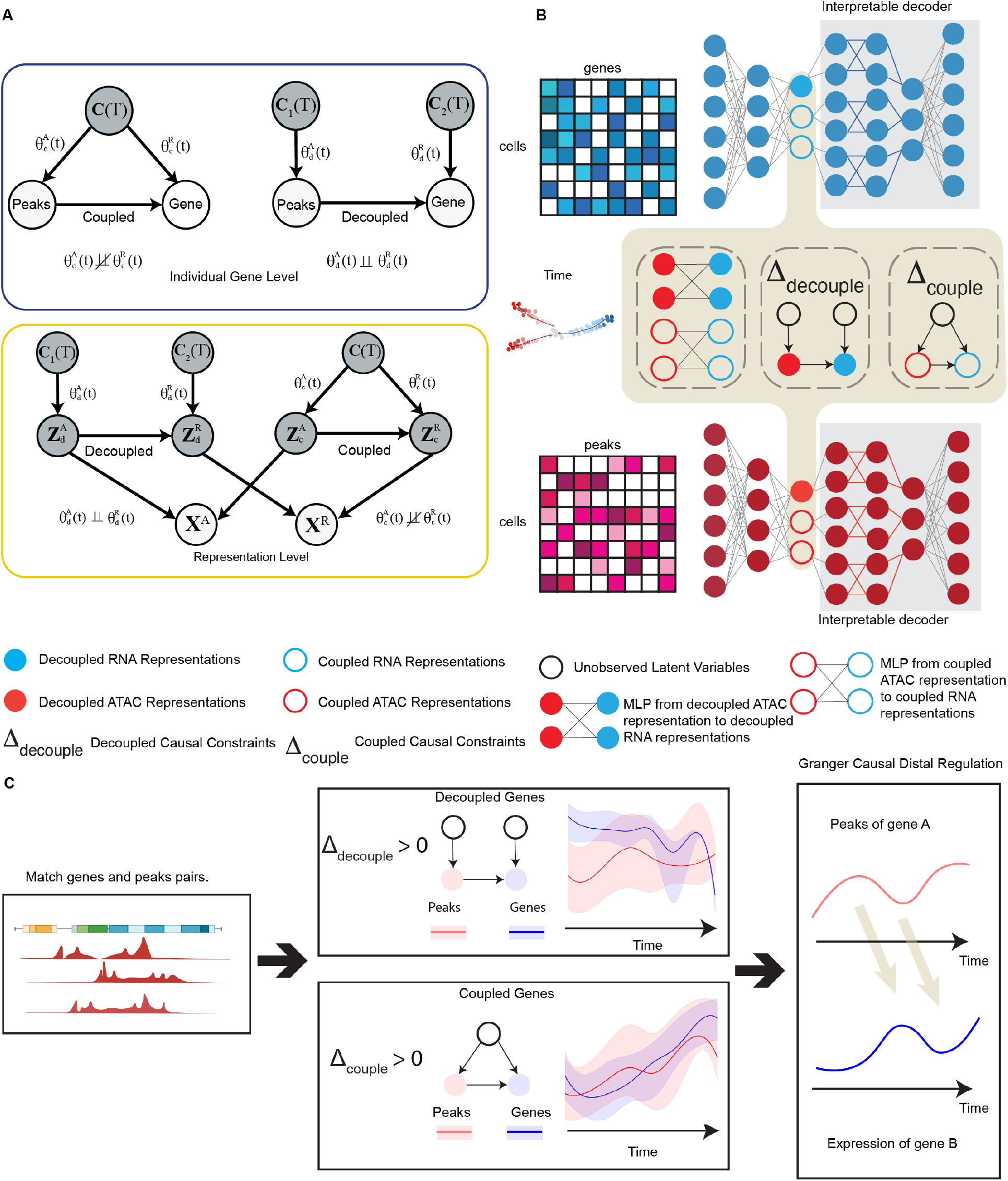
The Main Framework of HALO. **A**. Top: The causal diagram of individual gene expression and its corresponding peaks; Bottom: the causal diagram of representation level of scRNA-seq and scATAC-seq. **B**. Depicts the architecture for representation learning within a causal regularized variational autoencoder (VAE) framework. For the ATAC modality, the latent ***Z***^A^ is divided into 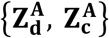, representing the decoupled and coupled latent representations, respectively. Similarly, the RNA modality’s latent representations comprise 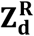 (decoupled) and 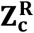 (coupled). The decoupled representations 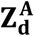 and 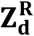 adhere to the decoupled causal constraints Δ_**decouple**_, whereas the coupled representations, 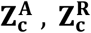 conform to the coupled causal constraints Δ_**couple**_. **C**. Illustrates the gene-peak level analysis process. Initially, genes and ATAC peaks within gene proximities are matched using non-negative binomial regression, linking gene expression to neighboring ATAC peaks. Subsequently, this analysis computes decouple and couple scores to categorize gene-peaks into either decoupled or coupled genes. Finally, the analysis employs the Granger causality test to identify distal regulatory relationships between peaks and genes, uncovering potential mechanisms of genetic regulation.

On the representation level, we model the lower dimensional latent space interactions between ATAC and RNA modalities from a causal perspective. Specifically, HALO learns the representations that contain the “coupled” information between scATAC-seq 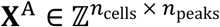 and scRNA-seq 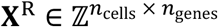 data as well as the independent “decoupled” information, such that the causal relations are preserved even across time (**Figure 1B**). Coupled representations encapsulate information where gene expression changes are dependent on chromatin accessibility over time, reflecting shared information across modalities. In contrast, decoupled representations extract information where gene expression changes independently of chromatin accessibility over time, emphasizing modality-specific information. The latent representations of scATAC-seq data can be decomposed to 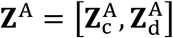. The latent representations of scRNA-seq data ***Z***^*R*^ may be similarly decomposed: 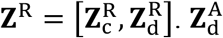 and 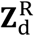 represent the decoupled latent representations derived from scATAC-seq and scRNA-seq data while 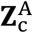 and 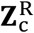 denote the coupled latent representations for scATAC-seq and scRNA-seq data, respectively. These representations capture distinct aspects of information from the two modalities, each varying over time or along a latent temporal variable. **Figure 1A** illustrates the underlying causal relationship, wherein 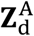 causes 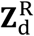, and 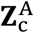 causes 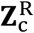. However, 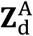 and 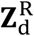 are influenced by independent causative factors, which are temporally related. This implies that, despite their causal linkage, 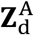 and 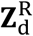 have independent causal factors, leading to independent temporal changes. In contrast to the decoupled representations, both 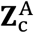 and 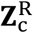 are influenced by common latent confounders, which also vary with time. This indicates that the changes in 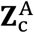 and 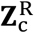 are synchronized over time, reflecting their shared temporal dynamics.

To ensure the delineation of these relationships, HALO utilizes paired scRNA-seq and scATAC-seq data as its input. We employ two distinct encoders to derive the latent representations ***Z***^A^ and ***Z***^R^. Moreover, we have formulated specific decoupled and coupled causal constraints (**see Methods Section 3)**. A Multi-Layer Perceptron (MLP) is used to model the concept that 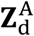 causes 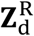, while a decoupled constraint enforces the independent functional relations between them. Similarly, another MLP is used to align the coupled representations 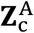 to 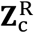, but with a coupled constraint (**See Methods Section 4**). Additionally, we developed a nonlinear interpretable decoder (**Figure 1B**) that allows us to interpret the latent representations by decomposing the reconstruction of genes or peaks into additive contributions from individual representations **(See Methods Section 5**).

At the individual gene level, our approach begins with the use of denoised gene expressions and peaks (**Figure 1C**), applying negative binomial regression to correlate local peaks with gene expression. This method allows us to match local peaks to corresponding gene expression enabling the subsequent calculation of decouple and couple scores at the individual gene level. Specifically, the decouple score quantifies the independence of gene expression level changes in relation to local peaks over time. Conversely, the couple score evaluates the extent to which gene expression changes are dependent on local peaks throughout the temporal course (**See Methods Section 6.3**). Finally, through the application of Granger causality analysis, we explore the underlying mechanisms of distal peak-gene regulatory interactions. This analytical approach allows us to elucidate instances where local peaks increase, yet corresponding gene expression remains largely unchanged.

In summary, HALO offers several key contributions to the field of multi-omics analysis, which we enumerate as follows: (1) HALO learns latent representations that are causally informed, enhancing our understanding of the interactions across different omics modalities. (2) HALO enables learning of interpretable latent representations. (3) HALO causally characterizes the relations between gene expression and associated chromatin regions over time, unlocking insights about regulatory dynamics. (4) HALO identifies distal *cis*-regulation interactions between chromatin regions and nearby genes, specifically for chromatin regions overlapping with super enhancers.

### HALO effectively separates coupled and decoupled representations, enhancing the analysis and interpretation of modality-shared and modality-specific information in mouse skin hair follicle data

Through the integration of information from both ATAC-seq and RNA-seq modalities, HALO is able to discern cell types and capture latent temporal dynamics. The ATAC and RNA representations from HALO are concatenated and visualized according to cell type (**Figure 2A, Figure S1A and S2**) and latent time (**Figure S1C**). **Figure 2B and F** display the UMAP projections constructed by the coupled RNA and ATAC representations, 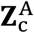 and 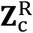, respectively. In accordance with the concept of coupled representations, 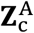 and 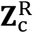 capture analogous information across the two modalities, which is reflected in the resemblance between their UMAP visualizations. However, the decoupled representations 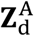 and 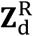 convey different information. In particular, 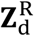 retains cell type information, while 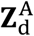 does not (**Figure 2C and G)**.

**Figure 2.**
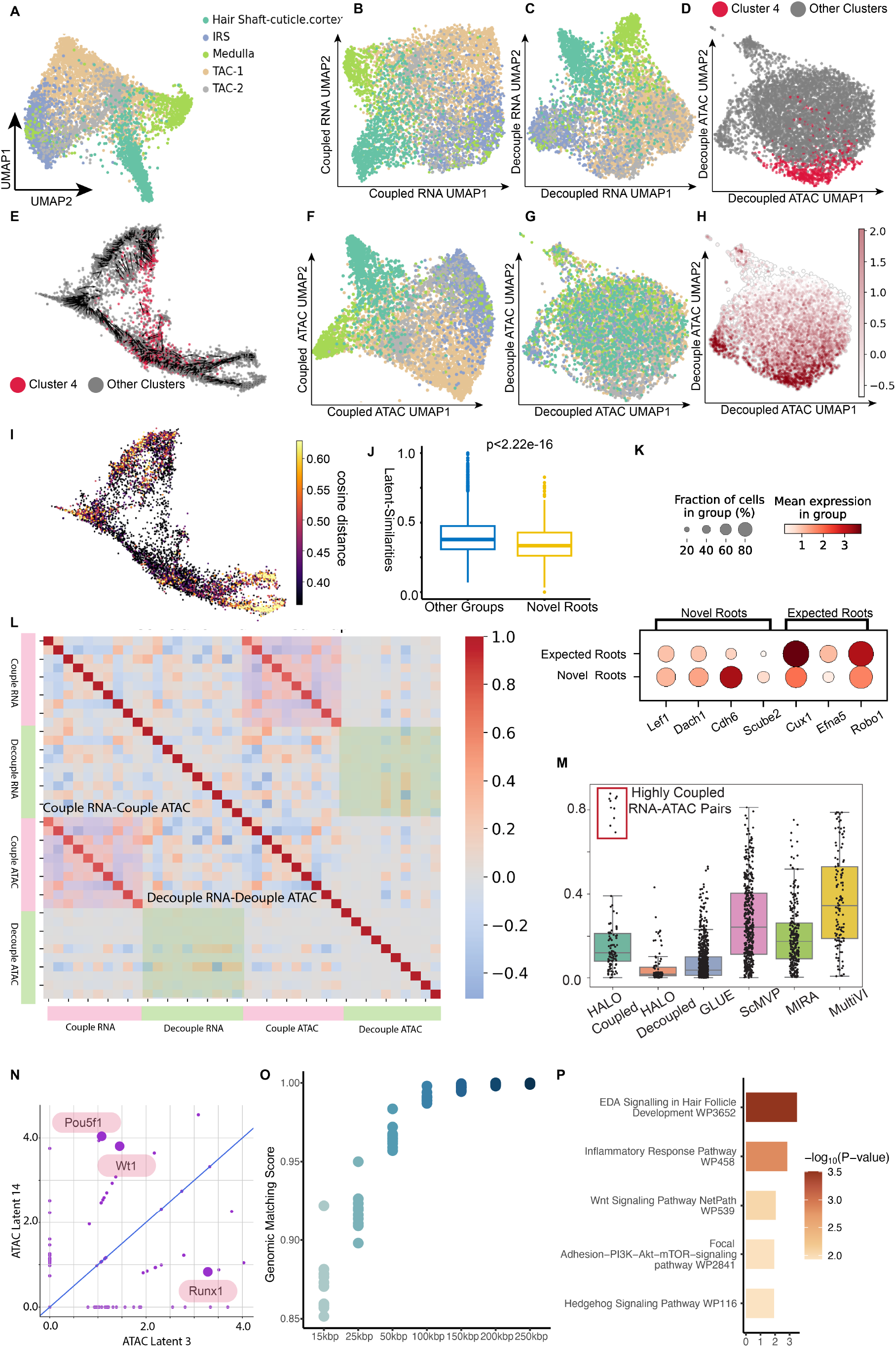
The representation level results of SHARE-seq mouse skin hair follicle dataset. The decouple representations contain the chromatin potential information in the latent space. **A**. The UMAP constructed from concatenated RNA and ATAC representations, colored by cell type. **B**. The UMAP constructed from RNA coupled representations, colored by cell type. **C**. The UMAP of RNA decoupled representations, colored by cell type. **D**. The UMAP constructed from ATAC decoupled representations, colored by cluster membership (cluster 4 and the rest of the clusters). **E**. The distribution of cluster 4 and other clusters, along with chromatin potential vector field, shown on previously published UMAP^8,46^. **F**. The UMAP constructed from ATAC coupled representations, colored by cell type. **G**. The UMAP constructed from ATAC decoupled representations, colored by cell type. **H**. The UMAP constructed from ATAC decoupled representations, colored by the value of ATAC decoupled latent representation 14, which mainly characterizes cluster 4. **I**. The original UMAP of SHARE-seq data^8^ colored by the cosine distance of decoupled RNA and decoupled ATAC representations. **J**. The cosine similarity between decoupled RNA and decoupled ATAC representations of cluster 4 (Novel Roots) and rest of cells. **K**. The marker genes expression levels of the novel roots and expected roots. **L**. The Pearson correlations matrix of latent representations. **M**. The Pearson correlations for HALO (coupled representations), HALO (decoupled representations), GLUE, scMVP, MIRA, and MultiVI. **N**. The enriched transcription factors (TFs) for coupled ATAC latent representation 3 and decoupled ATAC latent representation 14. **O**. The Genomic Matching Score of coupled representations with different genomic distances from genes’ TSS site. **P**. The gene ontology (GO) enrichment of top genes of RNA coupled representation 3.

Examining the analysis of the decoupled ATAC representation 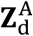, we performed clustering based on it. Cluster 4 (**Figure 2D)** is notably characterized by the decoupled ATAC latent representation 14 (**Figure 2H)**. Additionally, we have mapped cluster 4 onto the original SHARE-seq UMAP projection (**Figure 2E** and **Figure S1B**), revealing that cluster 4 corresponds to the novel root cells^8^. These novel roots cells were previously identified by chromatin potential, which is a quantitative measure of chromatin lineage-priming and used for cell fate prediction^8^. Moreover, it is also demonstrated that novel roots (cluster 4) express distinct marker genes in comparison to expected root cells (**Figure 2K)**. To validate this novel cell state in HALO’s framework, we evaluated the decoupled ATAC and RNA representations and found that, consistent with the definition of chromatin potential^8^, the cosine distance between the decoupled representations is maximal within the novel roots (**Figure 2I - J)**. By utilizing the interpretable decoder, HALO can identify the enriched transcription factors (TFs) within specific latent ATAC representations. **Figure 2N** shows the significantly enriched TFs in coupled representation 3 and decoupled representation 14, including Wt1 and Pou5fl. Transcription factors Wt1 and Pou5f1 are important in the Wnt/β-Catenin signaling pathway, exerting regulatory control over Lef1 and Dach1^9,10^, which serve as marker genes for novel root cells. In coupled ATAC representation 3, the transcription factor Runx1 is enriched, which directly promotes the proliferation of hair follicle stem cells and affects hair morphogenesis and differentiation. Coupled RNA representation 3, which is highly correlated with coupled ATAC representation 3, captures Eda, Wnt, and Sonic hedgehog (Shh) signaling pathways, which are important for hair follicle morphogenesis (**Figure 2P**)^11^.

To validate the relations between the coupled and decoupled representations of the two modalities, we show the Pearson correlation matrix for the latent representations (**Figure 2L**), which reveals a strong correlation between the ATAC and RNA coupled representations. In contrast, the decoupled representations exhibit a weaker correlation. Additionally, we assessed the correlations of latent representations between modalities using various representation learning methods, including MIRA^4^, GLUE^6^, MultiVI^5^, and scMVP^12^. The comparative analysis (**Figure 2M, S1D)** indicates that HALO’s decoupled representations maintain the lowest correlations, whereas its coupled representations are highly correlated across modalities with other four distinct datasets: Mouse Brain from 10X genomics, NEAT-Seq^13^, NeurIPS^14^ and Systemic sclerosis (SSc) pulmonary epithelium. We further interrogate whether the decoupled representations identified by HALO represent modality-specific batch bias or truly modality-specific biological information. To address this, we compared HALO’s performance in batch correction against that of other frameworks on the NeurIPS datasets^14^, which are known to contain substantial batch effects (**Figure S1E**), by silhouette score^15^ and Hilbert-Schmidt Independence Criterion (HSIC) (**see Methods Sections 8.2 & 8.3**). HALO is particularly effective at removing batch information, thereby confirming that its decoupled representations predominantly capture modality-specific biological information rather than mere batch effects.

To further examine our approach, we have developed a genomic matching score (**see Methods Section 8.1**) for coupled RNA and ATAC representation pairs that assesses the distance between their important genes and peaks by calculating the ratio of peaks that are located within the *cis*-regulation regions of the genes. **Figure 2O** presents genomic matching scores for the coupled ATAC-RNA representation pairs in the mouse skin hair follicle dataset, which demonstrates that HALO’s coupled representations could capture regulatory interactions between peaks and gene expressions from the extent to which they are aligned with each other.

### HALO characterizes gene-peak interactions in a temporal causal perspective

To further characterize the causal temporal relations between individual genes and peaks, we categorize gene-peak pairs into coupled genes and decoupled genes (**Figure 1A, 3B**). Employing negative binomial regression from denoised nearby peaks to denoised gene expression penalized by genomic distance, we align gene-peak pairs (**See Methods Section 6.1**). Upon aggregating these matched peaks, HALO calculates decouple and couple scores to quantitatively assess the extent of “decoupledness” and “coupledness” in gene-peak relationships, with positive scores suggesting decoupled and coupled, respectively. **Figure S3A-C** display the simulation outcomes for the decouple scores of both simulated decoupled and coupled gene-peaks pairs (**See Methods Section 11**). The SHARE-seq hair follicle data represent several lineages, differentiated from transit-amplifying cells (TACs) to inner root sheath (IRS), medulla, and cuticle/cortex cells (**Figure 3A**). **Figure 3C & D** and **S5 A & B** highlight the decoupled/coupled gene-peaks across all branches, with the respective couple and decouple scores (**Figure 3H, S4A & C**).

**Figure 3.**
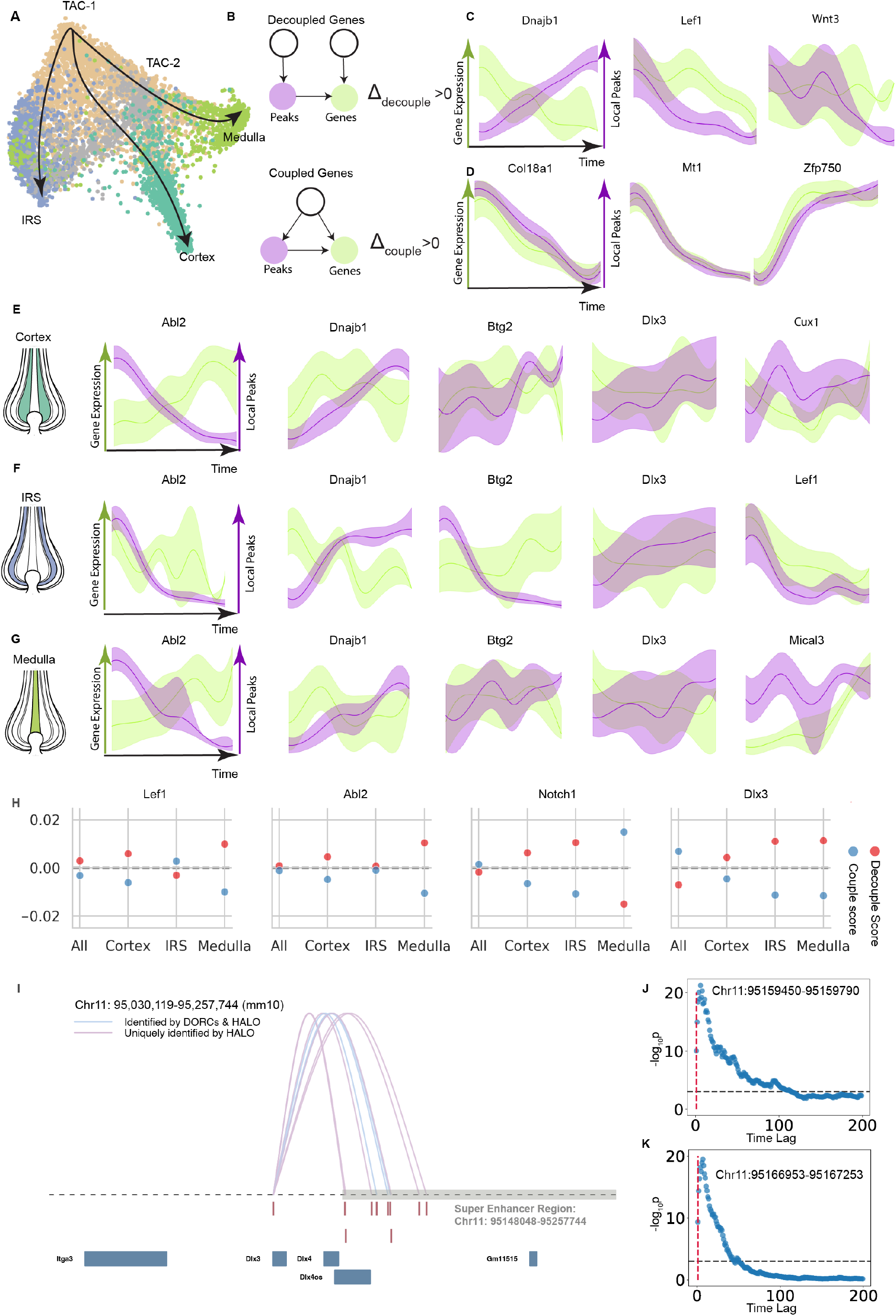
The individual gene level results of SHARE-seq mouse skin follicle hair dataset. **A**. The development trajectory of mouse hair follicle cells. **B**. The causal diagrams of decoupled and coupled individual gene-peak pairs. If the decouple score Δ_**decouple**_> 0, we classify the gene-peak pair as decoupled. If the couple score Δ_**couple**_> 0, we classify the gene-peak pair as coupled. **C**. Examples of decoupled (top) and coupled (bottom) gene-peak pairs across all branches. **D**. Examples of coupled gene-peak pairs across all branches. **E**. Examples of decoupled gene-peak pairs in Cortex. **F**. Examples of decoupled gene-peak pairs in inner root sheath (IRS). **G**. Examples of decoupled gene-peak pairs in Medulla. **H**. The decouple/couple score of genes on different developmental branches. **I**. The loops denote significant connections between peaks within super enhancer region and RNA expression of Dlx3. The connections between the expression of Dlx3 and peaks are identified by a gene-peak-linking algorithm and reported by original study^8^. **J**. The scatter plot visualizes significance of Granger causal relations between Itga3 gene expression and peaks in Chr11: 95159490-95159790. The X-axis is the time lag (number of cells, sorted by latent time), Y-axis is -log(P-values). Granger causality significance is evaluated by using the peaks of all previous cells [*c*_*t*−*n*_,*c*_*t*_) up to lag *n* (number of cells with index preceding time *t*) to determine the gene expression for cell *c*_t_, where *t* is determined by latent time sequential ordering of all cells. We utilize the likelihood ratio test for the Granger causality-based regulation inference. **K**. The scatter plot of Granger causal relations between Itga3 gene expression and peaks in Chr11: 95166953-95167253. The X-axis is the time lag (number of cells sorted by latent time), Y-axis is -log(P-values).

Additionally, HALO takes into account distinct developmental lineages: Cortex lineage (**Figure 3E, S5C & D**), IRS (**Figure 3F, S5E & F**), and Medulla (**Figure 3G, S5G & H**). By examining the decoupled genes (Abl2, Dnajb1, Dlx3 and Btg2) across different branches or within specific lineages, it becomes apparent that gene expression levels and corresponding peaks change independently over time. Specifically, Dlx3 exhibits decoupled behaviors on all three branches (**Figure 3H**); the gene expression remains relatively stable, but the aggregated peaks keep going up with time (**Figure 3E-G**). **Some genes (Notch1, Dlx3) exhibit coupled expression across all cells, but show decoupled behaviors within specific lineages, highlighting that gene regulation is highly context specific**.

We address the question of why these peaks continue to increase by examining the downstream effect of the accessibility of a local chromatin region on the gene expression of potential regulation targets other than the corresponding gene. Granger causality is well suited for the task of inferring these regulatory relationships due to the *time lag* between a local peak and target gene expression change. By utilizing the Granger causality test (**see Methods Section 10.1**), we are able to identify the distal regulation relations among Dlx3’s corresponding peaks and nearby genes other than Dlx3. In **Figure 3I**, the pink loops are peak-gene interactions inferred by DORC^8^, while the blue loops are uniquely detected by negative binomial regression. By predicting gene expression using the peaks of preceding cells with respect to ascending latent time, we find that nine local peaks within mouse skin hair follicle super enhancer region exhibit Granger causal relations with the expression of Itga3 (**Figure 3J-K** and **Figure S4B**)^16^. In summary, by analyzing gene-peak pairs at the individual gene level, HALO effectively distinguishes between coupled and decoupled interactions, as well as illustrates that local peaks of decoupled genes may contribute to expression of nearby genes.

### HALO uncovers regulatory factors in primary human CD4+ effector T cells assayed by NEAT-seq

NEAT-seq profiles the intra-nuclear protein epitope abundance of a panel T cell master transcription factors (TFs), chromatin accessibility, and transcriptome in single cells^13^. Due to posttranscriptional regulatory mechanisms affecting GATA3 protein expression^17^, we can leverage the protein quantification of the Th2 master TF GATA3 for more cell state information than can be gleaned from using RNA expression levels alone. HALO infers latent representations utilizing nuclear protein level of GATA3 as a proxy variable for latent time (**see Methods Section 7.2**) and constructs UMAP embedding to visualize distinct T cell subsets (**Figure 4A, S6 A-C, S7 & S8**). Notably, two coupled latent representations, RNA coupled 9 and ATAC coupled 9, show negative correlations with the nuclear protein level of GATA3 (**Figure 4 B-C**). RNA coupled 9 captures mTORC1 signaling (**Figure 4D**), which negatively regulates Th2 differentiation^18^. Consistently, ATAC coupled 9 is enriched in ZFX/NR4A motifs (**Figure 4E**), where NR4A is known to suppresses Th2 genes^19,20^. Conversely, Th17-specific ATAC coupled representation 12 exhibits enrichment in NR2F6/ARID5A motifs (**Figure S8 & 4E**), which are essential for the differentiation and function of Th17 cells^21,22^.

**Figure 4.**
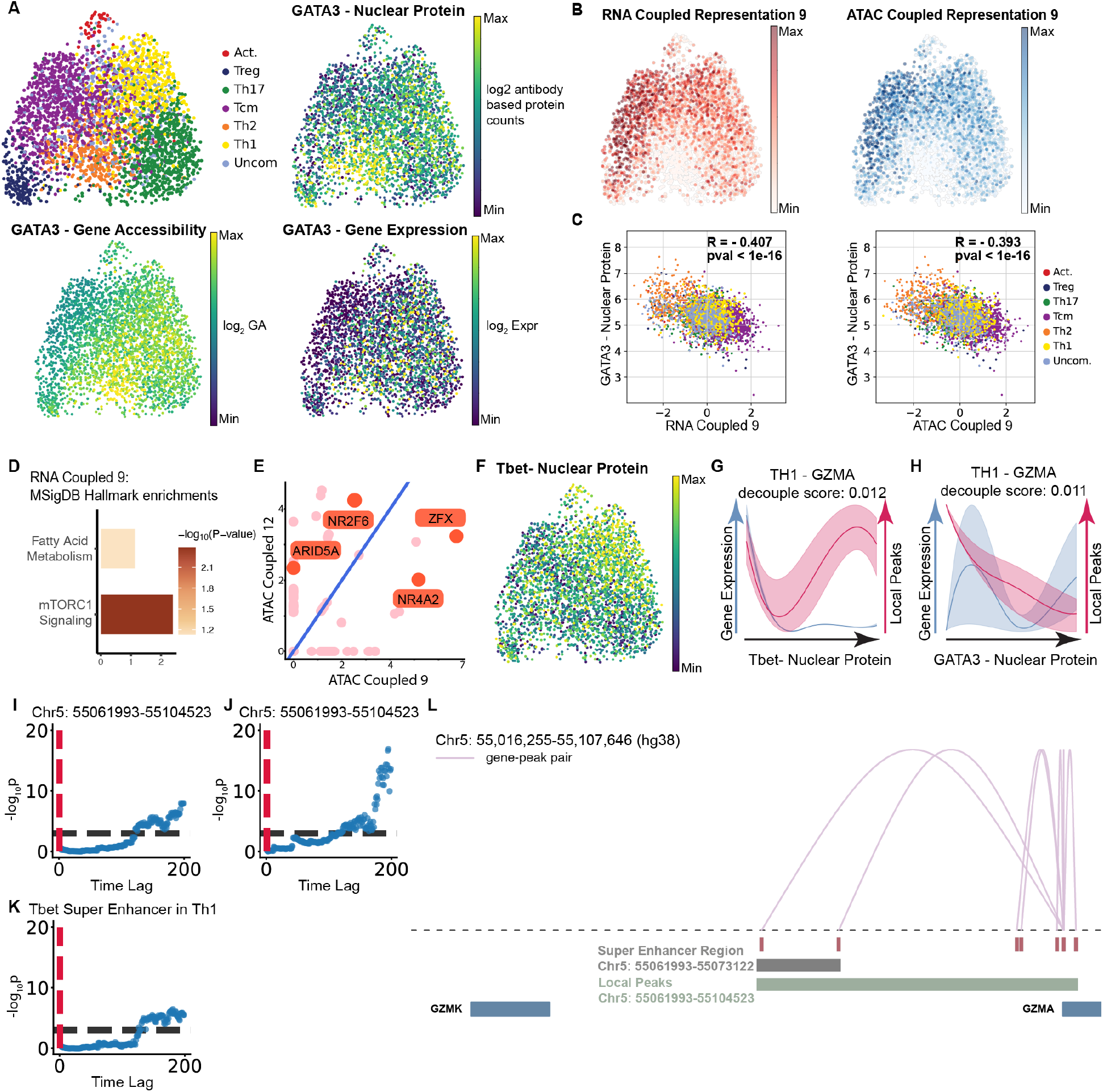
HALO reveals regulatory factors of human CD4+ effector T cell dataset. A. UMAP visualization of human CD4+ effector T cells constructed from concatenated RNA and ATAC representations, colored by original cell type, as well as nuclear protein, gene accessibility, and gene expression of GATA3. B. RNA coupled 9 and ATAC coupled 9 representations on UMAP embedding of RNA and ATAC representations. C. Scatter plots visualization GATA3 nuclear protein level and coupled latent representations (RNA coupled 9 and ATAC coupled 9) D. Gene set enrichment for RNA coupled 9. E. Motif enrichment for ATAC coupled 9 and ATAC coupled 12. F. Nuclear protein level of Tbet on UMAP embedding of RNA and ATAC representations. G. Gene-peak pairs of GZMA in Th1 cells sorted by Tbet nuclear protein level. H. Gene-peak pairs of GZMA in Th1 cells sorted by GATA3 nuclear protein level. I. The scatter plot of Granger causal relations between GZMK gene expression and peaks in Chr5: 55061993-55104523. The X-axis is the time lag (number of cells, sorted by Tbet nuclear protein level), Y-axis is -log(P-values). We utilized the F-test for the Granger causality-based regulation inference. J. The scatter plot of Granger causal relations between GZMK gene expression and peaks in Chr5: 55061993-55104523. The X-axis is the time lag (number of cells, sorted by GATA3 nuclear protein level), Y-axis is -log(P-values). K. The scatter plot of Granger causal relations between GZMK gene expression and peaks in Th1-specific T-bet super enhancer region (Chr5: 55061993-55073122). The X-axis is the time lag (number of cells, sorted by Tbet nuclear protein level), Y-axis is -log(P-values). L. The loops denote significant connections between local peaks and RNA expression of GZMA. The connections between the expression of GZMA and peaks are identified by a gene-peak-linking algorithm.

GATA3 regulates the Th2 cell fate decision in CD4+ T cells, while Tbet orchestrates Th1 differentiation. Both T-bet and GATA3 are co-expressed in Th1 cells, rather than being exclusively present in their respective lineages (**Figure 4F**). Within Th1 cells, T-bet suppresses Th2 genes by redistributing GATA3 from Th2 gene loci to Tbet binding sites within Th1 gene regions^23^. Granzyme genes (GZMA, GZMB, GZMK) possess binding sites specifically targeted by both T-bet and GATA3 in Th1 cells^24^. To dissect the regulation mechanisms of different TFs, we calculate decouple scores for Th1 cells using protein levels of GATA3 and T-bet as temporal information. Intriguingly, expression of the GZMA gene and its local peaks exhibit decoupled dynamics in Th1 cells (**Figure 4G-H**). As GATA3 protein levels rise, local peaks of GZMA become less accessible, even though GZMA gene expression is increased at high GATA3 protein levels (**Figure 4H**). Consistently, CD4+ T cells co-expressing T-bet and GATA3 exhibit upregulation of GZMA gene expression, compared to cells expressing only T-bet (**Figure S6D**). Conversely, increases in nuclear T-bet levels enhance the accessibility of local GZMA peaks, while GZMA gene expression level remains constant (**Figure 4G**). Using nuclear protein levels as temporal information, Granger causality tests reveal that local peaks of GZMA mediate the gene expression of GZMK (**Figure 4I-L**). Strikingly, local peaks of GZMA within the Th1-specific T-bet super enhancer region are also crucial for regulating gene expression of GZMK^25^. Leveraging TF protein levels as proxies for temporal information, HALO coupled representations not only exhibit strong correlation between RNA and ATAC modalities, but also capture similar biological contexts (**Figure S6E**). HALO allows us to further investigate the regulation of a TF using Granger causality test to determine downstream enhancer and its gene targets.

### HALO reveals epigenetic regulation of alveolar epithelial differentiation in systemic sclerosis-associated interstitial lung disease (SSc-ILD)

Systemic sclerosis (SSc) is an autoimmune disease characterized by fibrosis in the lungs and other organs. In SSc-ILD, alveolar epithelial cells are decreased due to impaired regeneration function, while airway epithelial cells (ciliated, club, goblet, basal) increase in fibrotic lesions^26^. Recent scRNA-seq analysis of human distal lungs and alveoli has shown that alveolar type-2 (AT2) cells acquire a unique epithelial transition state, known as AT0, during primate lung regeneration and disease^27^. AT0 cells have the potential to differentiate into either alveolar type-1 (AT1) cells or terminal and respiratory bronchiole-specific secretory cells (TRB-SCs) when cultured *in vitro*. However, in idiopathic pulmonary fibrosis (IPF), AT0 cells predominantly transform into terminal secretory cells within severe fibrotic areas, referred to as ‘bronchiolized regions’ in IPF lungs^27^.

To investigate the molecular underpinnings of impaired alveolar epithelial regeneration in SSc-ILD, we analyzed alveolar epithelial and terminal secretory cells using single-cell multiome sequencing data from six SSc-ILD and seven control lungs^28^. Due to its distinct genomic profile, airway basal cells were excluded from downstream analysis. We identified five clusters of alveolar epithelial and terminal secretory cells based on the expression of known marker genes (**Figure 5A, S9A-D**). Two subpopulations of AT2 cells (clusters 0 and 1) exhibit similar transcriptomic profiles but differ in chromatin accessibility. Additionally, we identified AT1 cells (cluster 3) and TRB-SCs (cluster 4), which co-express SFTPB and SCGB3A2, along with club and goblet cells (cluster 2).

**Figure 5.**
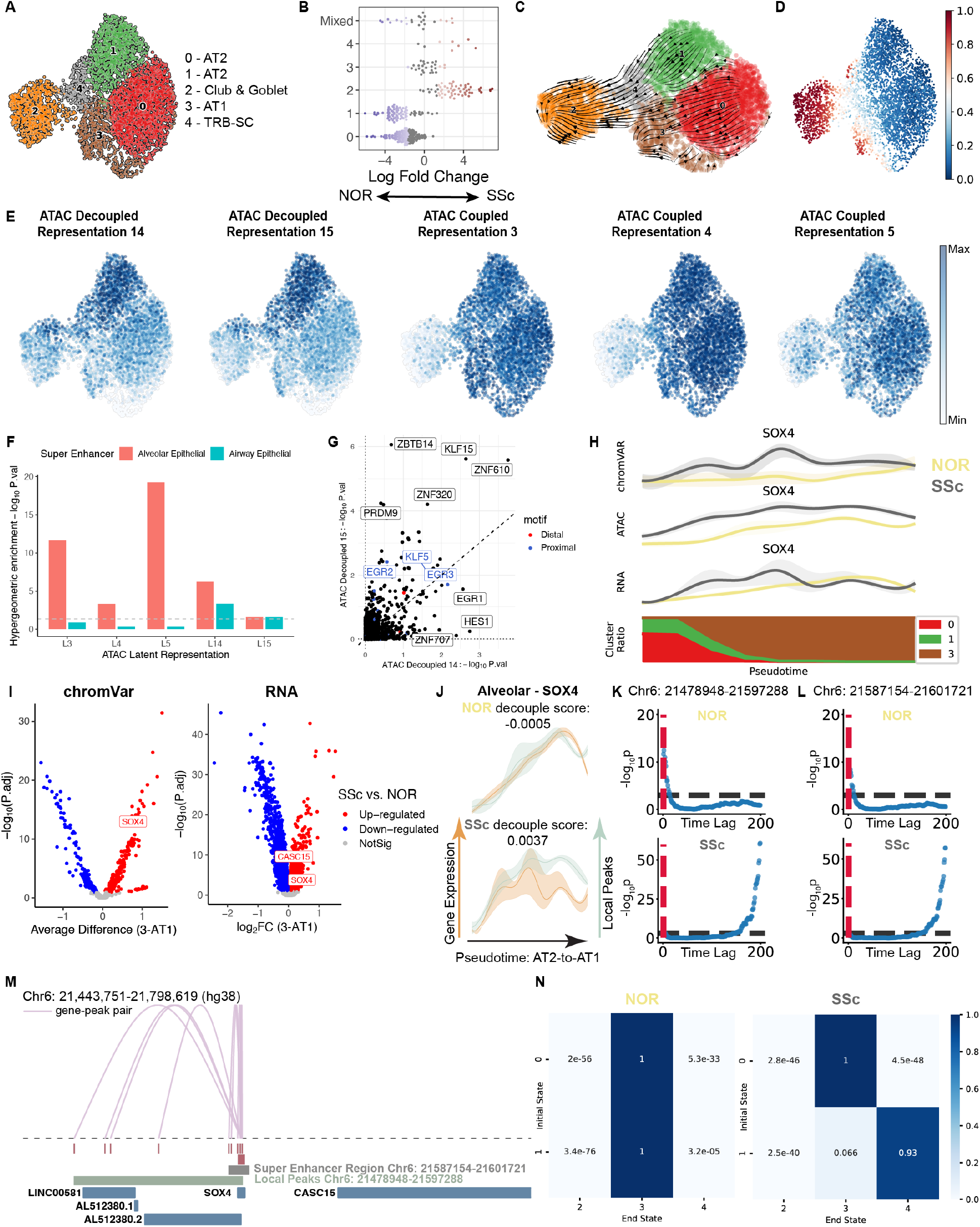
HALO uncovers chromatin lineage-priming in AT2 cells collected from SSc-ILD lungs. A. UMAP visualization of pulmonary epithelial cells constructed from concatenated RNA and ATAC representations, colored by identified cell types. B. Differential abundance test using Milo to compare control and SSc-ILD samples. C. RNA velocity analysis showing differentiation trajectories of pulmonary epithelial cells. D. UMAP visualization of pulmonary epithelial cells, colored by pseudotime. E. ATAC latent representations (3, 4, 5, 14, 15) on UMAP embedding. F. Hypergeometric enrichment of cell type specific super enhancer for ATAC latent representations. The dashed line indicates a p-value of 0.05. G. Motif enrichment for ATAC decoupled representations 14 and 15, where TFs essential for proximal and distal airway patterning during lung development are colored blue and red, respectively H. ChromVAR motif activity, local peaks, and gene expression of SOX4 in alveolar epithelial cells (clusters 0, 1, 3) from SSc and normal (NOR) lung, sorted by pseudotime. I. Volcano plots visualize SSc-associated changes in gene expression and ChromVAR motif activity in cluster 3 (AT1). J. Gene-peak pairs of SOX4 during AT2-to-AT1 differentiation (clusters 0, 1, 3) in alveolar epithelium, sorted by pseudotime. K. The scatter plot showing Granger causal relations of CASC15 gene expression and local peaks of SOX4 in Chr6: 21478948-21597288. The X-axis is the time lag (number of cells sorted by pseudotime), Y-axis is -log(P-values). We utilized the likelihood ratio test for the Granger causality-based regulation inference. L. The scatter plot showing Granger causal relations of CASC15 gene expression and AT1-specific super enhancer region (Chr6: 21587154-21601721). The X-axis is the time lag (number of cells sorted by pseudotime), Y-axis is -log(P-values). M. The loops denote significant connections between local peaks and RNA expression of SOX4. Connections between the expression of SOX4 and peaks are identified by a gene-peak-linking algorithm. N. Transition probability matrix of AT2 cell (clusters 0, 1) differentiation trajectories under different conditions, using optimal transport.

Our analysis revealed a substantial decrease in the AT2 cell population and an increase in secretory cells in SSc-ILD samples (**Figure 5B**). Additionally, AT2 cells in SSc-ILD lungs mirrored the transcriptional state of AT2 cells cultured in EGF-depleted organoids (**Figure S10A-E**), where AT2 cells convert to AT0 cells and subsequently progress to TRB-SCs following EGF depletion^27^. Consistently, trajectory inference algorithms (scVelo^29^, CellRank 2^30^, and Palantir^31^) applied to epithelial populations from both SSc-ILD and control samples reveal two terminal states in the AT2 differentiation trajectory. Cluster 1 progresses to secretory cells, passing through TRB-SCs, while cluster 0 differentiates into AT1 cells (**Figure 5C-D & S9E-F**).

To identify epigenetic regulations underlying AT2 lineage specification, we applied HALO to infer latent representations utilizing Palantir pseudotime. While RNA profiles alone are insufficient to distinguish the two AT2 subpopulations, decoupled ATAC representations are informative in separating these two subpopulations (**Figure S11, S12A-B**). Two decoupled ATAC representations (decoupled 14 and decoupled 15) characterize the transition from AT2 cells (cluster 1) to TRB-SCs (cluster 4) (**Figure 5E**). With an interpretable decoder, we find that top peaks contributing to the decoupled representations are enriched in airway epithelial-specific super enhancers and known transcription factors essential for proximal airway patterning during lung development^32,33^(**Figure 5E-G**). In contrast, three coupled ATAC representations describe the alveolar epithelial populations (cluster 0, 1, 3 & 4), which are enriched in alveolar epithelium-specific super enhancers (**Figure 5E-F**). Consistently, ChromVAR motif activity analysis indicates that cluster 1 exhibits lower motif activity for alveolar lineage transcription factors (NKX2-1 and CEBPA) compared to cluster 0, the other AT2 cluster (**Figure S10F**). CEBPA restricts AT2 cell plasticity during development and injury-repair, while CEBPA-dependent regulation recruits alveolar epithelial lineage TF NKX2-1 to promote and maintain the AT2 program^34–37^. Additionally, there are increased motif activities of transcription factors for proximal airway patterning and decreased motif activity of CEBPA in AT2 cells (cluster 0 & 1) from SSc lungs (**Figure S10G**). Analysis of modality-specific decoupled representations using HALO’s interpretable framework enables us to discover the epigenetic landscape and cell fates of alveolar epithelial cells.

We further evaluated transition probabilities among these clusters using optimal transport based on RNA and ATAC representations obtained from HALO (**see Methods Section 9**). In normal lungs, both AT2 clusters would progress to AT1. However, in SSc lungs, although cluster 0 still differentiated into AT1, cluster 1 predominantly transformed into TRB-SCs (**Figure 5N & S9G**), demonstrating that bipotent AT2 cells exhibit different cell fate decisions under SSc conditions. This suggests that HALO is able to depict a comprehensive portrait of alveolar differentiation, while defining epigenetic regulatory effectors of cell identity and pathology.

We then applied HALO to examine disease-associated decoupled genes. The epithelial-mesenchymal transition (EMT) master regulator SOX4 plays a crucial role in AT2-to-AT1 differentiation^38,39^. In SSc-ILD, both the expression and local peaks of SOX4 increase (**Figure 5H-I**). The dynamics of SOX4 gene expression and its local peaks become decoupled during terminal differentiation to AT1 in SSc lungs (**Figure 5J**); the local peaks remain accessible while the gene expression decreases. Strikingly, these local peaks of SOX4 are significantly Granger causally related to the expression of the EMT-associated long non-coding RNA (lncRNA) CASC15 (**Figure 5K-M)**. Additionally, local peaks of SOX4 within AT1-specific super enhancer regions contribute to the expression of CASC15. In SSc condition, regulatory events of SOX4’s local peaks exhibit longer time lag compared to the normal condition. A previous study has shown that CASC15 is upregulated in aberrant basaloid cells, an epithelial cell type displaying a partial EMT phenotype in ILD lungs^40^. Using HALO, we uncovered disease-specific decoupled genes that may contribute to SSc-ILD pathogenesis.

To further characterize dysregulated gene regulatory network of AT2 cells under SSc conditions, we inferred TF-CRE-gene linkages (**see Methods Section 14**). Among these, the EGR, RFX, TCF, and NFI family transcription factors (TFs) exhibited the largest positive average differences in chromVAR motif activity (**Figure S12C, Supp Table 1**). Notably, RFX family TFs are crucial for airway epithelial differentiation, while TCF and NFI family TFs play essential roles in alveolar epithelial differentiation, survival and regeneration^41–43^.

## Methods

See the Supplementary Methods Section.

## Discussion

The relationship between chromatin accessibility and gene expression is complex and often asynchronous. Previous methods assume that the two are synced with respect to time, missing nuances like time-lagging effects and independent regulatory mechanisms. HALO is the first to address these gaps by differentiating between coupled (dependent) and decoupled (independent) changes from a causal perspective. The framework operates at both representations and individual gene levels to learn interpretable modality-specific and shared information, as well as characterizes gene and associated peak interactions.

We conduct extensive benchmarks to evaluate the performance of HALO from different perspectives. Across several datasets, we demonstrate that HALO effectively learns highly coupled information across modalities while preserving decoupled and modality-specific information rather than batch bias. The highly coupled RNA and ATAC representation pairs, characterized by high Pearson correlation, capture functionally analogous biological contexts. In contrast, decoupled ATAC representations can inform our knowledge of chromatin accessibility-mediated cell fate potentials during the differentiation of mouse skin hair follicles and alveolar epithelium. Notably, we confirmed the bipotency of a subset of AT2 cells that acquire a previously reported transitional cellular state with the potential to differentiate into TRB-SCs or AT1 cells depending on niche signals^27,44^. We demonstrate that epigenetic information can shape this cell fate decision by conducting motif enrichment and cell type-specific super enhancer analyses on HALO’s decoupled representations. Achieving a deeper understanding of the factors underlying the diverging differentiation paths of AT2 cells opens the door to therapeutic options for a variety of lung diseases.

At the individual gene level, HALO distinguishes between coupled and decoupled gene-peak pairs. For decoupled cases where associated chromatin regions become more accessible, but gene expression remains stable or decreases, HALO employs Granger causality test to assess distal regulatory interactions between associated peaks and nearby genes. We find that peaks overlapping with super enhancer regions often exhibit such decoupled behavior and are involved in distal regulation during cellular transition or developmental trajectories. Additionally, HALO uncovers SSc-ILD-specific regulatory mechanisms, such as the decoupling of SOX4 expression and its local peaks during AT2-to-AT1 differentiation. We show that during this differentiation under SSc conditions, chromatin regions in the intersection of SOX4 local peaks and AT1-specific super enhancers are responsible for regulation of the lncRNA CASC15.

To enhance HALO’s applicability, future work can extend the framework to include profiling of additional modalities, such as methylation and protein levels, to provide a more comprehensive understanding of gene regulation dynamics. Additionally, leveraging spatial epigenome-transcriptome co-profiling technologies could allow HALO to uncover spatiotemporal dynamics and genome-wide gene regulation mechanisms within tissue contexts^45^. These advancements would extend HALO’s utility as a valuable tool for understanding the functions and regulatory mechanisms of cell populations.

## Appendix: HALO Methods

### 1 Table of Symbols and Variables

**Table 1.**
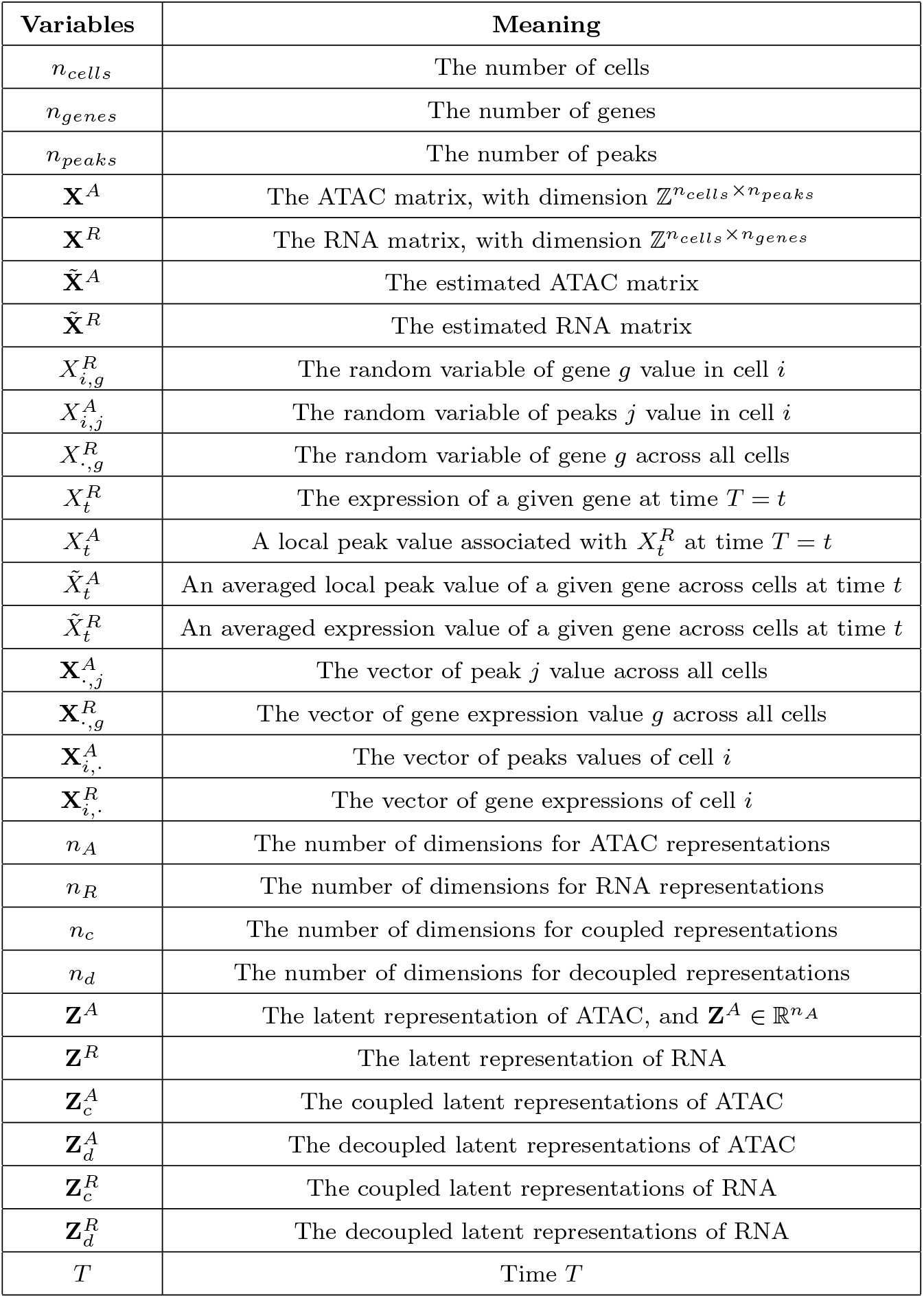
Table of Symbols and Variables.

| Variables | Meaning |
| --- | --- |
| $n_{cells}$ | The number of cells |
| $n_{genes}$ | The number of genes |
| $n_{peaks}$ | The number of peaks |
| $\mathbf{X}^A$ | The ATAC matrix, with dimension $\mathbb{Z}^{n_{cells} \times n_{peaks}}$ |
| $\mathbf{X}^R$ | The RNA matrix, with dimension $\mathbb{Z}^{n_{cells} \times n_{genes}}$ |
| $\tilde{\mathbf{X}}^A$ | The estimated ATAC matrix |
| $\tilde{\mathbf{X}}^R$ | The estimated RNA matrix |
| $X_{i,g}^R$ | The random variable of gene $g$ value in cell $i$ |
| $X_{i,j}^A$ | The random variable of peaks $j$ value in cell $i$ |
| $X_{\cdot,g}^R$ | The random variable of gene $g$ across all cells |
| $X_t^R$ | The expression of a given gene at time $T = t$ |
| $X_t^A$ | A local peak value associated with $X_t^R$ at time $T = t$ |
| $\tilde{X}_t^A$ | An averaged local peak value of a given gene across cells at time $t$ |
| $\tilde{X}_t^R$ | An averaged expression value of a given gene across cells at time $t$ |
| $\mathbf{X}_{\cdot,j}^A$ | The vector of peak $j$ value across all cells |
| $\mathbf{X}_{\cdot,g}^R$ | The vector of gene expression value $g$ across all cells |
| $\mathbf{X}_{i,\cdot}^A$ | The vector of peaks values of cell $i$ |
| $\mathbf{X}_{i,\cdot}^R$ | The vector of gene expressions of cell $i$ |
| $n_A$ | The number of dimensions for ATAC representations |
| $n_R$ | The number of dimensions for RNA representations |
| $n_c$ | The number of dimensions for coupled representations |
| $n_d$ | The number of dimensions for decoupled representations |
| $\mathbf{Z}^A$ | The latent representation of ATAC, and $\mathbf{Z}^A \in \mathbb{R}^{n_A}$ |
| $\mathbf{Z}^R$ | The latent representation of RNA |
| $\mathbf{Z}_c^A$ | The coupled latent representations of ATAC |
| $\mathbf{Z}_d^A$ | The decoupled latent representations of ATAC |
| $\mathbf{Z}_c^R$ | The coupled latent representations of RNA |
| $\mathbf{Z}_d^R$ | The decoupled latent representations of RNA |
| $T$ | Time $T$ |

**Table 2.**
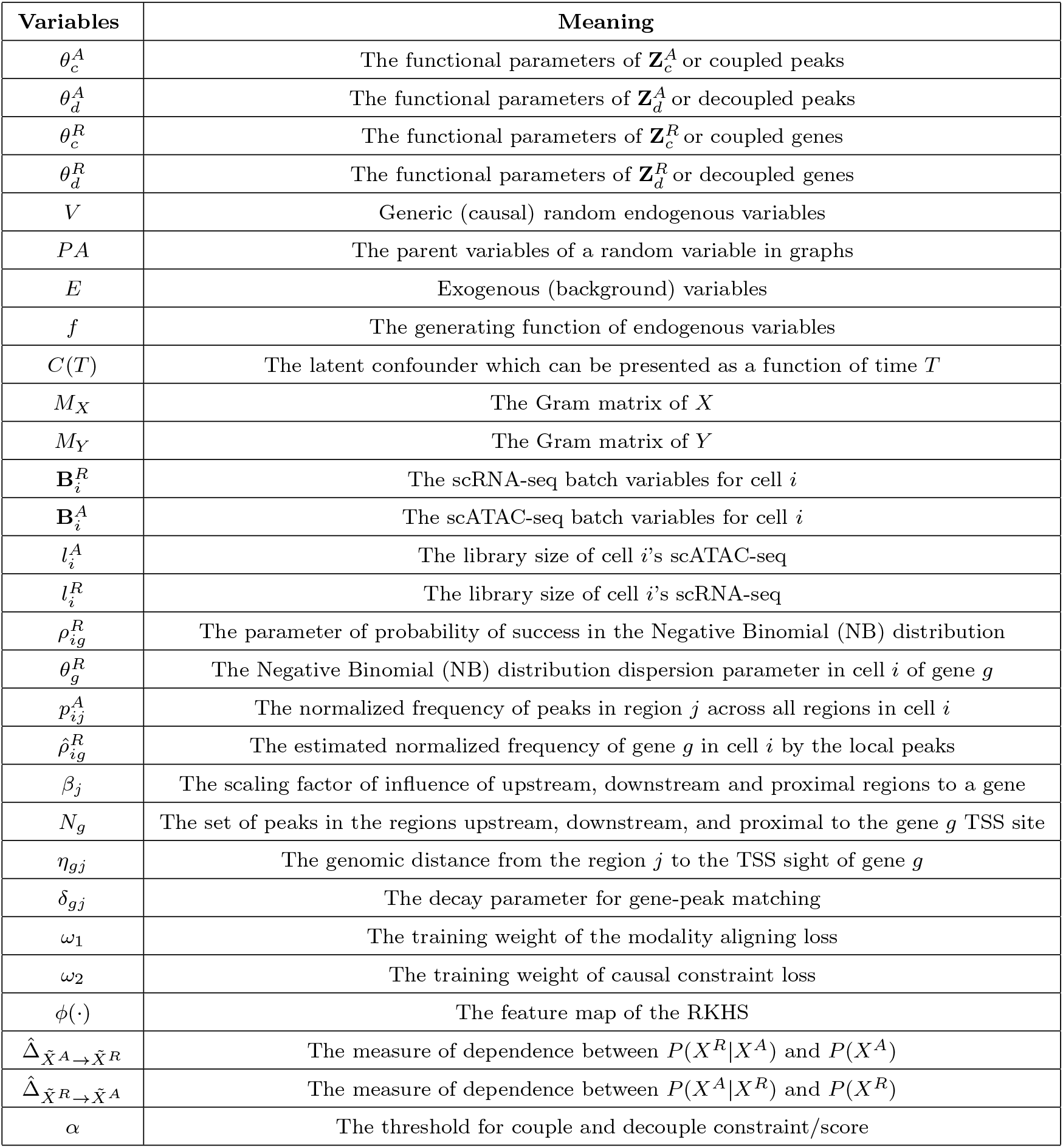
Table of Symbols and Variables, continued.

### 2 Problem Definition

In this section, we give the formal definition of our model. The input is co-assayed scATAC-seq and scRNA-seq data. Specifically, scATAC-seq 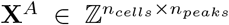 is a matrix with *n*_*cells*_ cells and *n*_*peaks*_ peaks, scRNA-seq 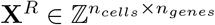 is a matrix with *n*_*cells*_ cells and *n*_*genes*_ genes. Moreover, 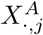 represents the value of peak *j* across all cells, 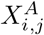 represents the value of peak *j* in cell *i*. 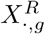 is the count for gene *g* across all cells, 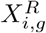 is the expression of gene *g* in cell *i*.

The low-dimensional latent causal representations of scATAC-seq data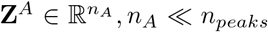 can be decomposed to 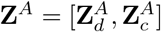, where 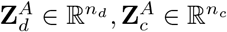, and *n*_*c*_ + *n*_*d*_ = *n*_*A*_. The latent representation of scRNA-seq data 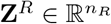 may be similarly decomposed: 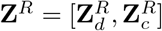, where 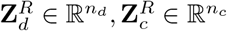, and *n*_*c*_+*n*_*d*_ = *n*_*R*_. For the sake of simplicity, we set *n*_*c*_ = *n*_*d*_ and 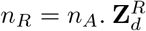 and 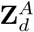 denote the decoupled latent representations of scRNA- and scATAC-seq data, while 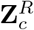 and 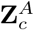 are their coupled counterparts. Moreover, following the general case [1], we denote parameters of the structural causal model (SCM) for coupled and decoupled latent representations as the following: 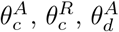, and 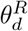 are the functional parameters of 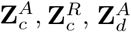, and 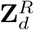, respectively. To model the dynamics/changes of the representations with time, we further assume that these functional parameters can be written as functions of time *T*, that is 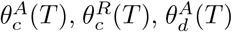, and 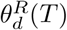. For simplicity of notation, we omit the *T* in the remainder of this paper.

**Appendix Fig. 1:**
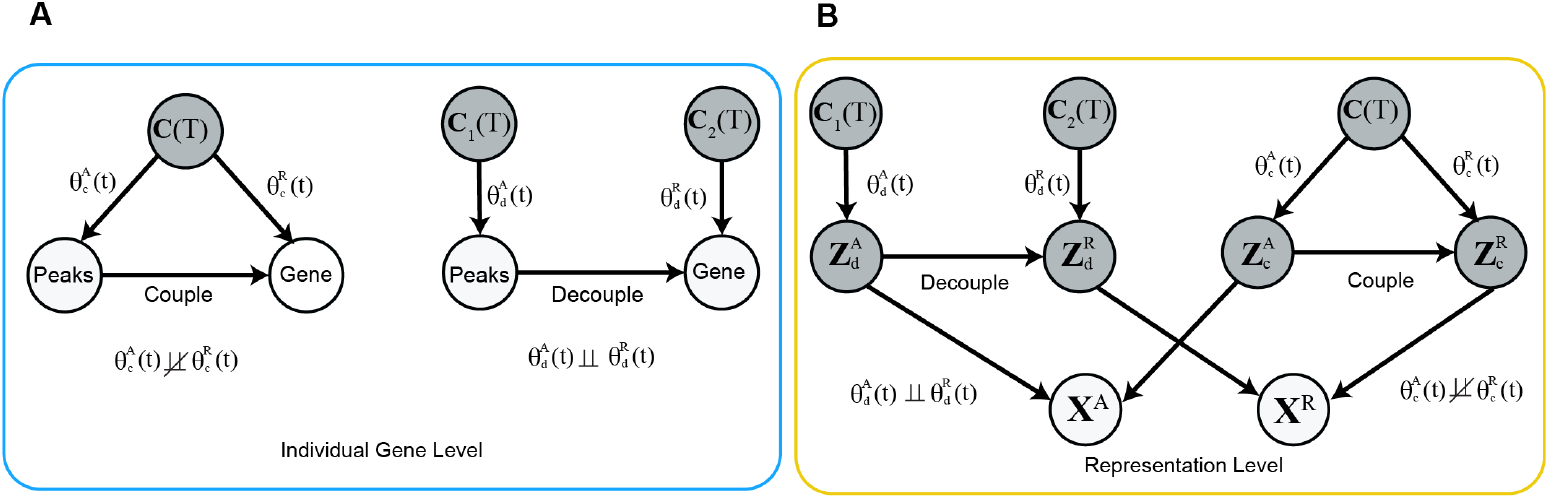
The causal diagram of the scATAC-seq and scRNA-seq. A. The causal diagram at individual gene level. The decoupled gene-peak pairs have independently changing parameters while coupled gene-peak pairs have dependently changing parameters. B. The causal diagram at representation level. Coupled and decoupled latent representations are denoted as scATAC-seq 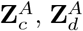, and scRNA-seq 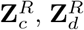. Similarly, coupled representations have dependently changing parameters, while decoupled representations have independently changing mechanisms.

### 3. Causal Constraints

In this section, we briefly introduce causal graphic models, structural causal models (SCMs), and non-stationary causal discovery conditions, then we introduce our causal constraints on representations.

#### 3.1 Causal Graphic Models and Structural Causal Models

Finding underlying causal relations is a fundamental task in many disciplines [2]. Moreover, utilizing causal models to encode distributional and causal assumptions is becoming increasingly popular [3–5]. Markovian causal models are one of the simplest model types and consist of a directed acyclic graph (DAG) over a set of vertices *V* = {*V*_1_,. .., *V*_*n*_}, representing variables of interest, as well as a set of directed edges, or arrows, that connect these vertices. The interpretation of such a graph has two components: probabilistic and causal. The probabilistic interpretation treats the arrows as representations of probabilistic dependencies among the corresponding variables, and the missing arrows represent conditional independencies. Each variable is independent of all its nondescendants given its direct parents in the graph [2, 6]. These assumptions are represented in the joint probability *P* (*V*) = *P* (*V*_1_,. .., *V*_*n*_) factorization as the following product:

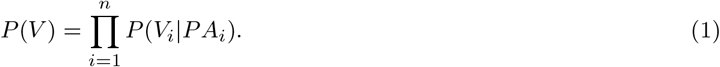

Here, *P* (*V*_*i*_ |*P A*_*i*_) is the conditional distribution of *V*_*i*_ given its parents *PA*_*i*_. The causal perspective of these models views the arrows as representing causal influences between the corresponding variables. In this case, the factorization in Eq. (1) still holds, but the factors are further assumed to represent autonomous data generation processes (here we refer to each *P*(*V*_*i*_|*PA*_*i*_) as a causal module or causal mechanism [1]); that is, each conditional probability *P*(*V*_*i*_|*PA*_*i*_) represents a stochastic process by which the values of V_*i*_ are generated in response to the values of *PA*_*i*_, and the stochastic variation of this assignment is assumed to be independent of the variations in all other assignments. In this sense, this representation is an alternative to an SCM. An SCM is defined [6] as a triple *M* = ⟨*E, V, F*⟩, where:

1. *E* is a set {*E*_1_, *E*_2_,. .., *E*_*n*_} of exogenous (background) variables that are determined by factors outside the model;
2. V is a set {*V*_1_, *V*_2_,. .., *V*_*n*_} of endogenous variables that are determined by variables in the model, *E* ⋃ *V* ; and
3. *F* is a set of functions {*f*_1_, *f*_2_,. .., *f*_*n*_} such that each *f*_*i*_ is a mapping from *E*_*i*_ ⋃ *P A*_*i*_ to *V*_*i*_, where *E*_*i*_ ⊆ *E* and *P A*_*i*_ ⊆ *V* \*V*_*i*_. That is, *V*_*i*_ = *f*_*i*_(*P A*_*i*_, *E*_*i*_), for *i* = 1,. .., *n*, assigns a value to *V*_*i*_ that is dependent on a set of variables in *V* ∪ *E*.

#### 3.2 Coupled and Decoupled Latent Representations

In most biological processes, the biological mechanisms change with time, and this displacement is referred to as *distribution shift*. If there are distribution shifts (i.e. *P* (*V*) changes across domains or over time), there must also be changes in the causal modules *P* (*V*_*i*_ | *P A*_*i*_),*i* ∈ *N* ; these are called *changing causal modules*. In the setting of our framework, this corresponds to changes over time in the mechanism by which chromatin accessibility affects gene expression. It is usually assumed that the quantities that change over time can be written as functions of a time index, *T* [1]. We assume *pseudo causal sufficiency*, which means that any confounders can be written as smooth functions of time. Hence, in our multi-omic cases, the latent confounders can be presented as a function of pseudo-time. Denoting **C**(*T*) as the set of pseudo confounders (this set may also be empty), we can write the SCM as:

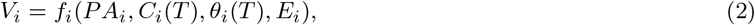

where *C*_*i*_(*T*) ⊆ **C**(*T*) denotes the set of confounders related to *T* which influence *V*_*i*_, θ_*i*_(*T*) denotes the effective parameters in the model that are also assumed to be functions of *T*, and *E*_*i*_ is the exogenous variables that is independent of *T* and *PA*_*i*_, with a non-zero variance.

We construct a multi-modal representation learning model through a causal lens, incorporating temporal dynamics. Drawing from prior knowledge, we recognize that certain parts of scATAC-seq and scRNA-seq data exhibit strong correlations over time, whereas others do not (Appendix Fig. 1A). Our causal modeling framework incorporates this temporal information and the core hypothesis that open chromatin causes active transcription. Specifically, we pose the causal relations as 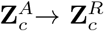 and 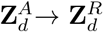.

We incorporate low dimensional representations in this biological process. Moreover,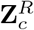 and 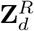 alto-gether generate scRNA-seq, while decoupled and coupled representations 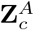 and 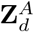 altogether generate scATAC-seq (Appendix Fig. 1B). Additionally, the independently changing mechanisms can be modeled as independently changing parameters of the generating functions. The correlated changing mechanisms can be formulated as confounded parameters of the generating functions. To be specific, it can be shown to be the following:

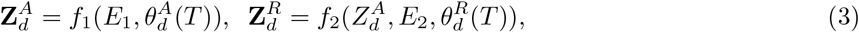

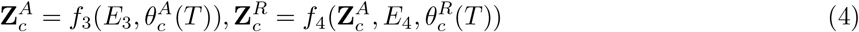

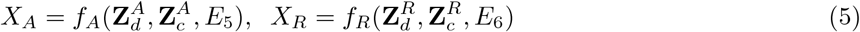

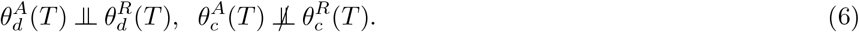

Further, we have following joint distributions,

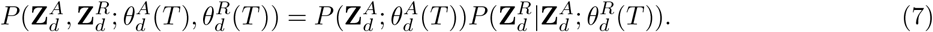

#### 3.3 Hilbert Schmidt Independence Criterion (HSIC)

Testing for the independence of changing *P* (**cause**) and *P* (**effect** |**cause**) can be achieved by using Hilbert Schmidt Independence Criterion with properly designed kernel embeddings of nonstationary conditional distributions [1]. Given a set of observational samples *{*(*x*_1_, *y*_1_), (*x*_2_, *y*_2_), …, (*x*_*N*_, *y*_*N*_)*}* for random variables *X* and *Y*, HSIC is a statistic for testing statistical independence by evaluating the squared covariances *C*_*XY*_ between feature maps of *X* and feature maps of *Y* . Let us introduce the following theorem:

##### Theorem 1.

*[7] Denote by ℱ, 𝒢 Reproducing Kernel Hilbert Spaces (RKHS) with universal kernels k, l on the compact domains 𝒳 and 𝒴, respectively. We assume without loss of generality that* ∥ *f* ∥_∞_ ≤ 1 *and* ∥ *g* ∥_∞_ ≤ 1, *for all f 2 F and g 2 G. Then* ∥ *C*_*xy*_ ∥ _*HS*_ = 0 *if and only if x* ∈ 𝒳 *and y* ∈ 𝒴 *are independent*.

Specifically, let **M**_*X*_ and **M**_*Y*_ be the Gram matrices for *X* and *Y* estimated on the observed samples. We use following estimator of HSIC [8]:

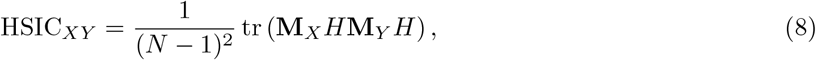

where *H* is for centering the features, with entries 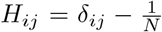, and *6*_*ij*_ = 1 when *i* = *j*. Moreover, the normalized HSIC is given by:

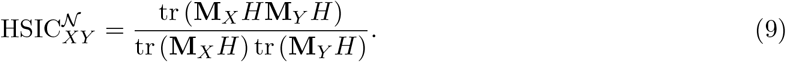

HSIC can be tailored to serve as a powerful tool for evaluating the dependence score between changing causal modules. However, instead of directly applying it to the scATAC-seq and scRNA-seq data, which tend to be noisy and sparse, we extend this approach to the latent space representations of these two modalities. In the next section, we introduce the formal definition of our problem and its solutions.

#### 3.4 Independently Changing Constraints on Representations

We are now ready to introduce the causal constraints on the representations. We postpone the topic of determining the time information *T* of the sequencing data to the Time Estimation section. For now, let us assume we have time information and further develop statistical constraints for the coupled and decoupled causal representations. As presented in the previous section, the joint distribution in the decoupled case can be formulated as such:

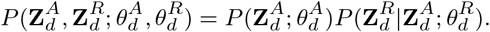

Based on this factorization, in light of CD-NOD [1], we formalize the specific joint distribution 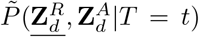 and its embeddings. 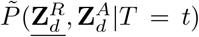 is designed to capture the changes in only the conditional distribution 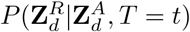. Formally, it is defined as

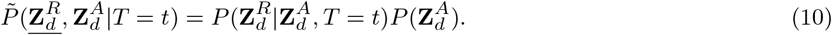

Note that in Eq. (10), 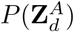 does not depend on *T* . As a result, 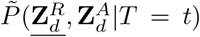 encodes only the changes across *T* in 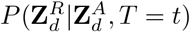. A corresponding joint distribution can be also applied to the coupled latent space. Let **Z**^*R*^ *2 Ƶ*^*R*^ and (ℋ, *k*) be a Reproducing Kernel Hilbert Space (RKHS) with a measurable kernel on *Ƶ*^*R*^. Let 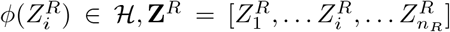 be the feature map for **Z**^*R*^ *2 Ƶ* ^*ℛ*^, and assume 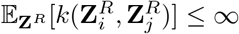. Similar notations are used for **Z**^*A*^, 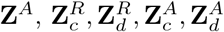, and *T* . Thus the general embedding of the joint distribution of coupled or decoupled latent representations can be formulated as the following:

##### Proposition 2.

*[1] Let* **Z**^*A*^ *represent the direct causes of* **Z**^*R*^, *and suppose that they have N observations. The kernel embedding of distribution* 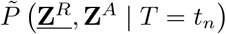 *can be represented as*

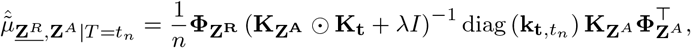

*where* 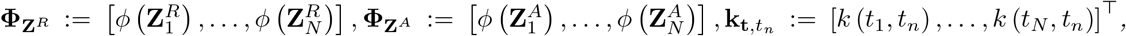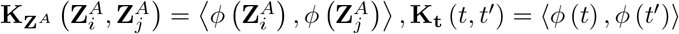 *and ⊙ represents point-wise product*.

The proof of this proposition can be found in the CD-NOD paper [1].

Our goal is to characterize the changing causal mechanisms between the latent spaces of scATAC- and scRNA-seq. Here we focus on the causal constraint for the representation level; for the gene-level analysis, please refer to Section Gene-Peak Matching. So for the decoupled case, 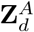 and 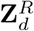 have independently changing mechanisms along *T* within the cell lifespan, but 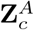 and 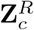 change dependently. Thus, we aim to enforce that 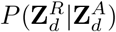 and 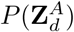 change independently, and that 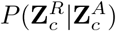 and 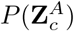 change dependently. CD-NOD [1] proposes a score function based on HSIC which measures the dependence of the changes in the two distributions *P* (*Y* |*X*) and *P* (*X*) over time. We translate the two-variable case to our **Z**^*A*^ and **Z**^*R*^ setting as follows:

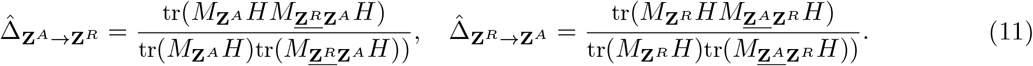

where 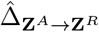 measures the dependence between *P* (**Z**^*R*^|**Z**^*A*^) and *P* (**Z**^*A*^). Similarly, 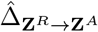 measures the dependence between *P* (**Z**^*A*^|**Z**^*R*^) and *P* (**Z**^*R*^). Between these two dependence measures, if the value of 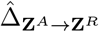 is smaller, we posit that *P* (**Z**^*R*^|**Z**^*A*^) and *P* (**Z**^*A*^) change independently, and that the causal relation is **Z**^*A*^ → **Z**^*R*^. In our case, for the decoupled representations 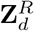 and 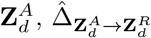 should be smaller than 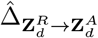, based on our knowledge of chromatin accessibility and transcription. On the other hand, for the coupled case, 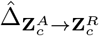 and 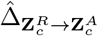 should be similar in value and both larger than a threshold α [1].

Formally, the causal constraints of 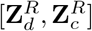 and 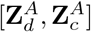 are given by the constraints as follows,

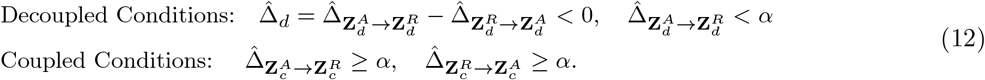

### 4 Generative Models

We introduce our generative model designed to learn the lower dimensional representations 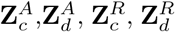. Utilizing inputs from scRNA-seq **X**^*R*^ and scATAC-seq **X**^*A*^, along with modality-specific batch information **B**^*R*^ and **B**^*A*^, we use mean field variational inference to estimate 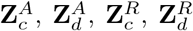 [9, 10]. To model the expression value of gene *g* in cell *i*, denoted 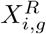, we specify the following distributions.

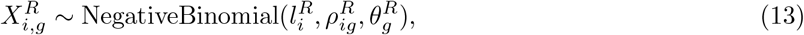

where 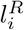 is the scaling factor in cell *i*, 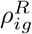 is the normalized composition of gene *g* across all genes in cell *i*, and 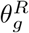 is the gene dispersion. Similarly, for chromatin region *j* of cell *i*, we consider 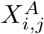 as a Multinomial distribution,

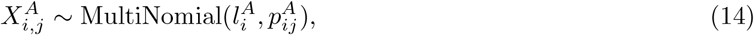

where 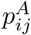 is the normalized frequency of accessibility across all regions in each cell, 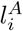 is the library size (scaling factor) for cell *i*. Moreover, we use 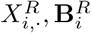 and 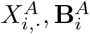 to infer library sizes 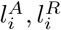 of each cell by variational inference, where 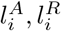 follow log-norm distributions, 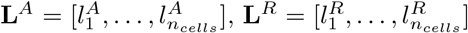 are the vectors of the library size of gene expression and peak of all cells. The mean field variational is used to estimate the latent representations with the following factorized posterior:

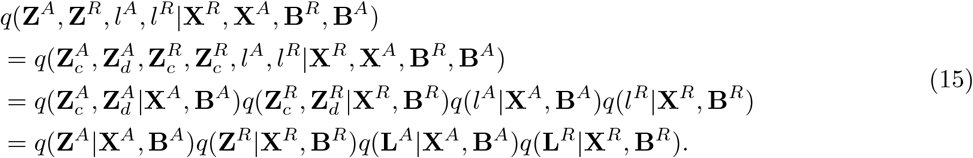

Then we use the following Evidence Lower Bound (ELBO),

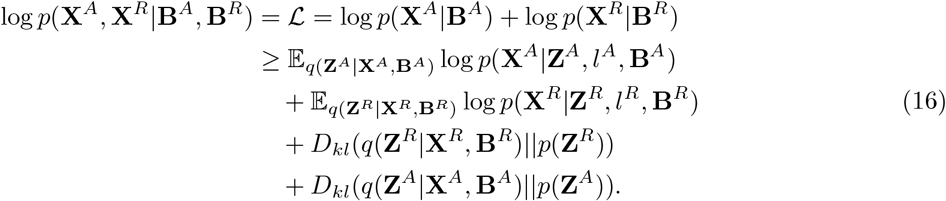

Specifically, we have the following posterior distributions,

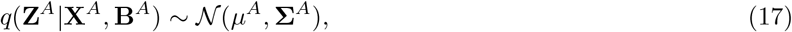

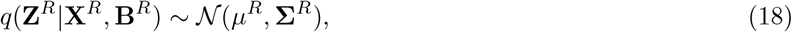

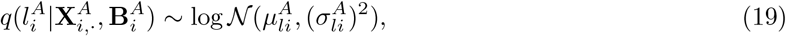

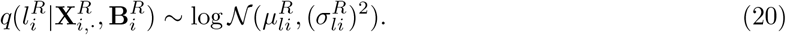

where *µ*_*A*_, *µ*_*R*_ are the means and Σ_*A*_, *Σ*_*R*_ are the covariance matrices of **Z**^*A*^, **Z**^*R*^, respectively. We use deep neural networks to estimate the following:

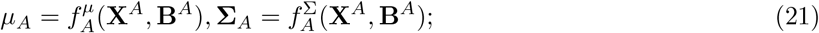

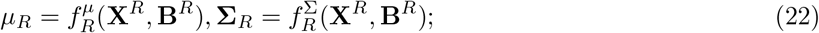

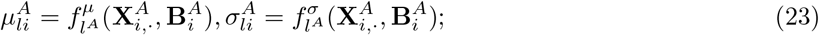

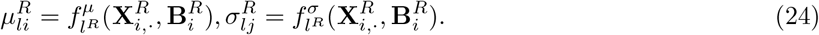

The prior distribution is specified as standard Gaussian distribution,

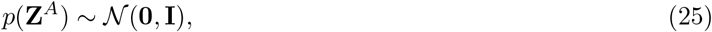

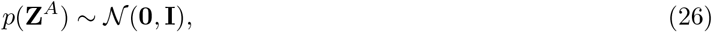

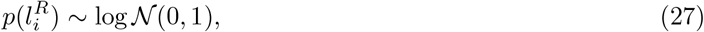

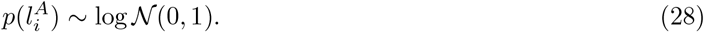

Next, we integrate the causal constraints outlined in the previous section into our final optimization objective. Both the coupled and decoupled representations are subject to the constraints specified in Equation (12). Thus, we define the causal constraint 𝒞 with a hyperparameter α,

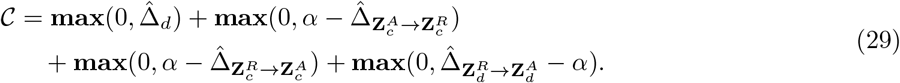

Moreover, we want to align the paired decoupled and coupled representations so that under the coupled or decoupled assumptions, 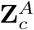 causes 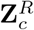 and 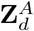 causes 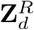. To realize these assumptions, we add the following layers,

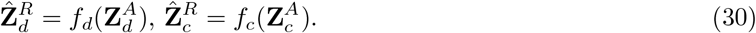

We want to minimize the *L*2 norm, ℳ:

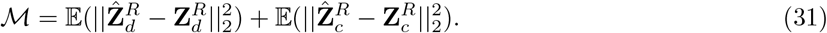

The main loss can be defined with parameters *ω*_1_, *ω*_2_ as follows:

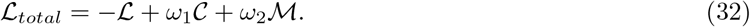

The ablation study of hyperparameters α, *ω*_1_, *ω*_2_ can be found in section Hyperparameters Tuning.

Thus the optimization objective is as follows,

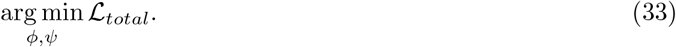

### 5 Neural Interpretable Decoder

We introduce the neural interpretable decoder in the HALO framework. In light of Neural Additive Models [11], for each latent variable, we utilize a sub-neural network with non-linear transformations to capture information specific to each individual latent variable. A logistic regression layer is added to the end of the neural network (Appendix Fig. 2). Thus, by analyzing the logistic regression weights (loading matrix), we can conclude how much one latent variable *Z*_*i*_ for *i* ∈ [1,. .., *n*] contributes to the reconstruction of a specific gene or peak (*X*). The non-linear transformation of latent representations can be formulated as follows,

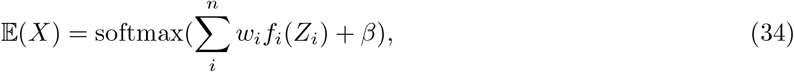

where *f*_*i*_ for *i* ∈ [1,. .., *n*] are the non-linear functions which can be presented as neural networks, *w*_*i*_ are the weights of the the last linear layer, and β is the bias of the last layer. Moreover, *f*_*i*_ is constructed as follows:

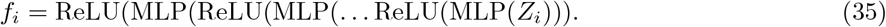

Since all *f*_*i*_ for *i* ∈ [1,. .., *n*] are non-negative, the weights *w*_*i*_ can interpreted as the importance/contribution of *Z*_*i*_ for the reconstruction of *X*.

#### 5.1 Important Genes and Peaks for Representations

For each representation, we are able extract the important genes and peaks to further interpret the latent representations. Specifically, for 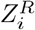, the *i*th element of the latent representations, we sort genes by the weights of last linear layer of the neural network, and pick the top 100 genes as the most “significant” genes for 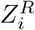. Similarly, we are able to retrieve the top 100 peaks as the most “significant” peaks for 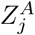.

**Appendix Fig. 2:**
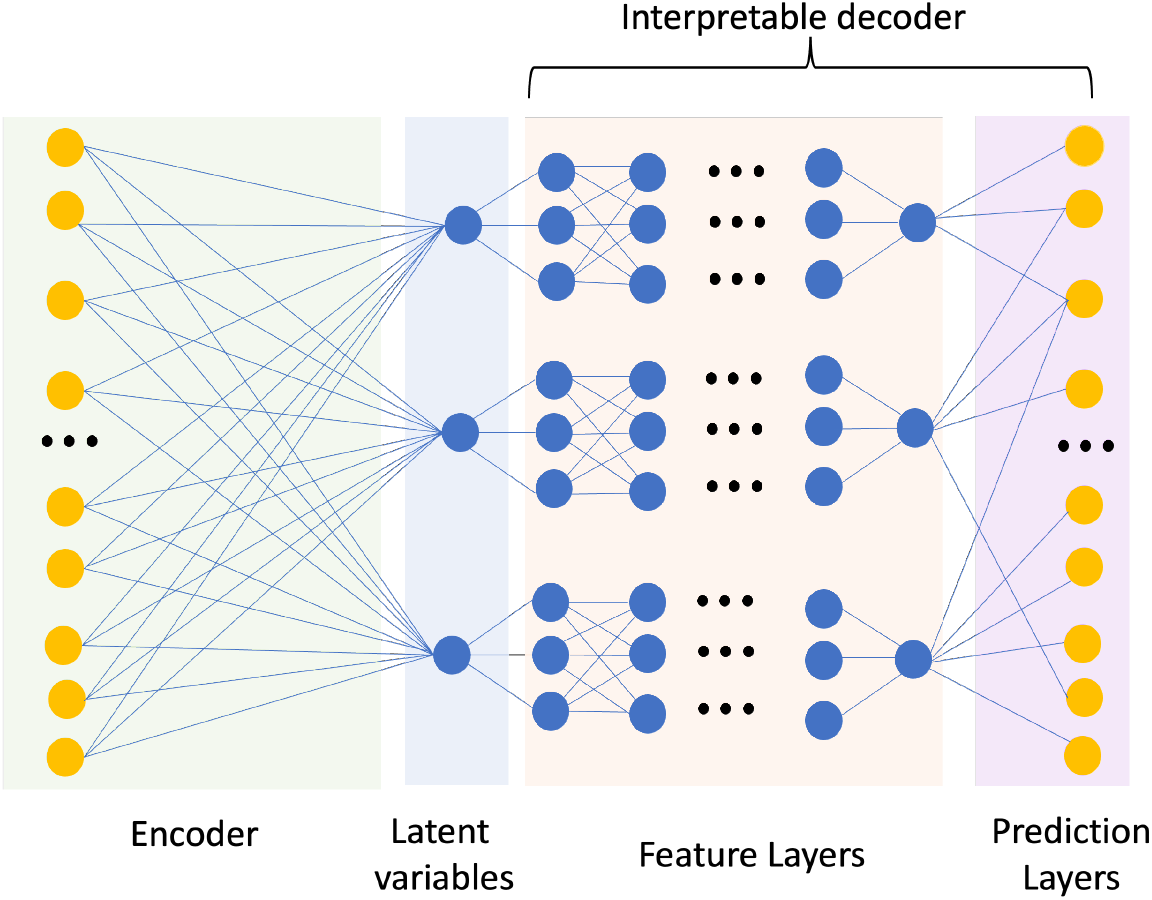
Structure of the interpretable decoder

### 6 Individual Gene Level Analysis

#### 6.1 Gene-Peak Matching

In light of MIRA [12], HALO correlates local scATAC-seq peaks to gene expression values, by searching within a certain distance upstream and downstream of local peaks to maximize the probability of observing gene expression values. Specifically, for gene *g ∈* {1, *n*_*genes*_} in cell *i*, as well as estimated chromatin accessibility probability at chromatin region *j* and its probability 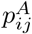 (see Section Generative Models), we have the following definitions:

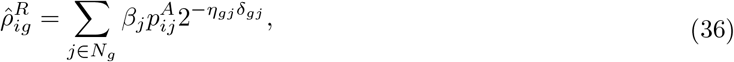

where *N*_*g*_ is the set of peaks *j* in the regions upstream, downstream, and proximal to the TSS of gene *g*; β_*j*_ ∼ HalfNormal(0, 1) is the scaling factor for the influence of peaks in upstream, downstream, and proximal regions to the gene *g*; η_*gj*_ is the TSS distance; and δ_*gj*_ is the decay parameter. HALO learns these parameters {β_*j*_, *η*_*gj*_, δ_*gj*_} by maximizing the likelihood of estimated 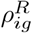 in the previous representation learning phase. The upstream and downstream region sets encompass regions between 1.5 and 600 kbp from the TSS, while the proximal region is within 1.5 kbp. HALO estimates the parameters by maximizing the scaled 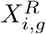 likelihood:

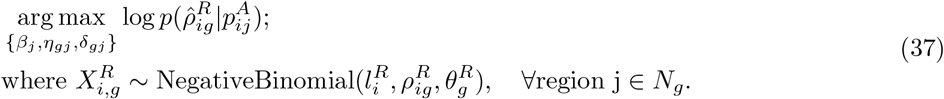

The optimization procedures follow MIRA [12].

#### 6.2 Individual Gene Causal Conditions

In this section, we apply the causal constraint to the individual gene level. After gene-peak pair matching, we can calculate the gene-level decouple score, further depicting the gene-peak behavior by characterizing how individual gene and corresponding matched peaks change with time. Recall the definition of coupled and decoupled causal relations: given time *T*, coupled gene-peak pairs have shared latent confounders that cause them to change in a correlated manner; in contrast, the decoupled gene-peak pairs have independently changing mechanisms with time, but are still associated in a causal manner. Appendix Fig. 1A illustrates the coupled and decoupled gene-peak relations. As described in previous sections, we can calculate the independence of the parameters of peaks and genes to characterize the degree of their “decoupledness.”

Similarly, given a gene *g*, its expression value *X*^*R*^, and associated peaks *X*^*A*^ (for simplicity of notation, we drop the gene and cell indices here). we define the individual gene level changing joint distribution as follows:

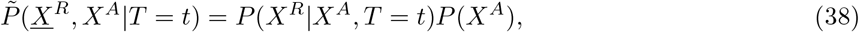

where 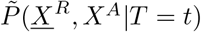 captures the changes of *X*^*R*^ with *T* that are not introduced by *X*^*A*^. Then we can calculate its kernel embeddings according to the proposition below. Let *X*^*R*^ ∈ *𝒳*^*R*^ and (ℋ, *k*) be a Reproducing Kernel Hilbert Space (RKHS) with a measurable kernel on *𝒳*^*R*^. Moreover, 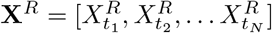, and 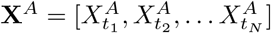 are samples we have for gene expression and associated peaks, where *t*_1_,. .., *t*_*N*_ are the time labels for each sample and *N* is the number of samples. Let ϕ (*X*^*R*^) ∈ *ℋ* be the feature map and assume 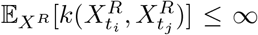 and 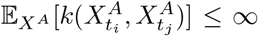. Thus the general embedding of the joint distribution of coupled or decoupled latent representations can be formulated as the following:

##### Proposition 3.

*Let X*^*A*^ *represent the gene g’s corresponding peaks which are the direct causes of the expression value X*^*R*^ *of gene g, and suppose that they have N observations. The kernel embedding of distribution* 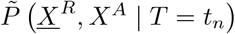 *can be represented as*

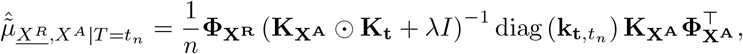

*where* 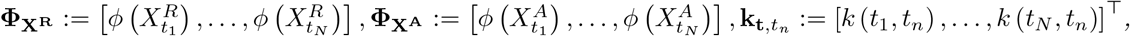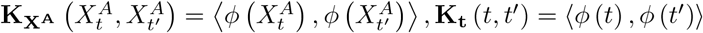 *and ⨀ represents point-wise product*.

With this proposition, we define 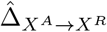 to measure the dependence between *P* (*X*^*R*^|*X*^*A*^) and *P* (*X*^*A*^), 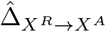 to measure the dependence between *P* (*X*^*A*^|*X*^*R*^) and *P* (*X*^*R*^),

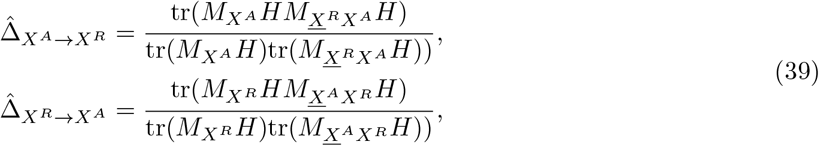

where 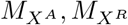 are the Gram Matrices [13] for *X*^*A*^ and *X*^*R*^, *H* is used to center the feature with entries 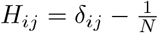, and δ_*ij*_ = 1 when *i* = *j*.

Calculating 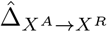 and 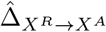 is challenging because single cell sequencing data suffer from sparsity problems. We utilize the aggregated estimated peaks 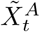 as 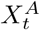 and averaged gene expression 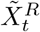 as 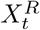 for all cells with time label *t*. Specifically,

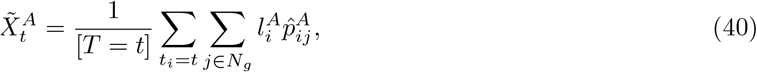

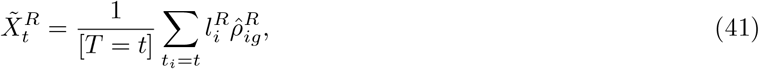

where *N*_*g*_ is all matched peaks for gene *g* in the upstream, downstream, and proximal areas of the TSS (See section Gene-Peak Matching), [*T* = *t*] is the number of samples/cells where the time label is equal to 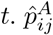 is the estimated parameter 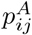 for peak *j*’s Multinomial distribution. 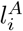 is the estimated library size for peaks (scaling factor), 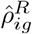 is the shape variable in cell *i* of gene *g*’s NB distribution, and 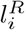 is the estimated library size for gene expression. Given this estimated 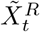 and 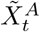, we can further calculate gene level couple and decouple scores.

#### 6.3 Gene Level Couple Score and Decouple Score

By utilizing the individual gene level causal constraints, we develop couple and decouple scores to measure the “coupledness” and “decoupledness” of genes and their associated peaks. Given the estimated aggregated gene *g* expression 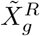, and its corresponding aggregated peaks *j* value 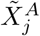. Note “average” gene and peaks mean that the gene or peaks are averaged across all cells or cells in a specific branch. To simplify notation, we drop the *i* and *j* in the rest of this section. Given a threshold value *α*, we define the following scores,

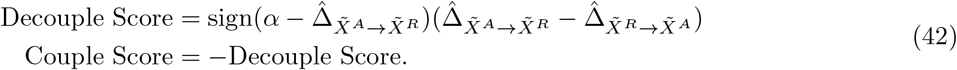

From the definition of decouple score and couple score, given a threshold value α, if the decouple score is positive, we can identify gene *g* and peaks *j* as decoupled; otherwise if the couple score is positive, *g* and peaks *j* are coupled.

### 7 Time Estimation

#### 7.1 Latent Time Estimation

Recall the need to address how to estimate the latent time and incorporate it into our framework. Generally speaking, we can use any latent time or pseudotime estimation methods to annotate cells. For the SSc human lung epithelium dataset, we use Palantir pseudotime to order cells [14], and for the mouse brain, mouse skin datasets, we utilize latent time inferred using MultiVelo [15].

#### 7.2 Protein-based Latent Time Estimation

We can also utilize the normalized levels of important intra-nuclear proteins as a proxy variable for latent time. In the NEAT-seq dataset of CD4 memory T cells, the transcription factor (TF) antibody-derived tag (ADT) count data were normalized using NPC antibody counts and hashtag oligos (HTOs), as described in the original study [16]. The NPC normalization can be formulated as,

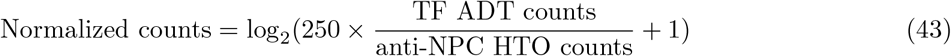

#### 7.3 METAcell Construction

If temporal information is unavailable, we can infer representations of paired scRNA-seq and scATAC-seq data based on reconstruction loss. Since cells at a similar developmental stage should have similar omics profiles, individual cells exhibiting similar states are identified then aggregated into METAcells [17, 18]. These METAcells are constructed using cell-cell similarities inferred from the latent representations, which account for potential decoupling phenomena between scRNA-seq and scATAC-seq data. We then assess distribution shifts by utilizing METAcell membership information *C*.

Here, we summarize the averaged peaks 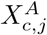 and averaged gene expression 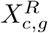 for all cells with METAcell membership label *c*. Specifically,

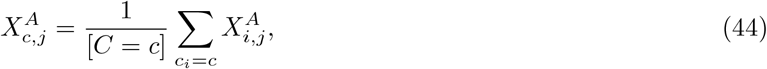

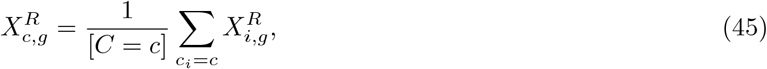

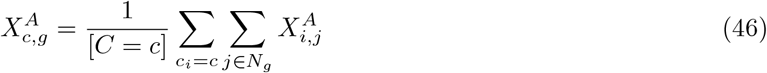

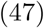

*c*_*i*_ represents the membership of cell *i*. [*C* = *c*] is the number of samples/cells assigned to a METAcell label *c*. Given these summarized values 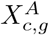 and 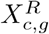, we can further calculate gene-level couple and decouple scores.

### 8 Evaluation Metrics

#### 8.1 Genomic Matching Score

To evaluate the distance between important genes and peaks from coupled RNA and ATAC representation pairs, we calculate the genomic matching score. This score assesses the fraction of important peaks within the ATAC representation that lie within the distal gene regulation distance (up to 250kbp) from the transcription start site (TSS) of significant genes in the RNA representation.

First, we select pairs of coupled RNA and ATAC representations with the highest Pearson correlation coefficients. Next, we identify significant RNA features and peak features that contribute to the latent representations. Specifically, for the SHARE-seq dataset, we extracted the top 100 RNA features and the top 2000 peak features from the corresponding coupled representation pairs.

#### 8.2 Silhouette Score

The Silhouette width [19] is used to evaluate the extent to which sample dissimilarity is minimized within a batch and maximized across batches. For a sample *x, d*_*a*_(*x*) is the average dissimilarity between *x* and all other data points of the batch *A* that *x* belongs to, while *d*_*c*_(*x*) is the average dissimilarity between *x* and the samples in batch *C* where *C ≠ A*, meaning that min(*d*_*c*_(*x*)) is the average dissimilarity for the batch that is most similar to *A*. Using these definitions, we have the following silhouette width *s*(*x*) of *x*,

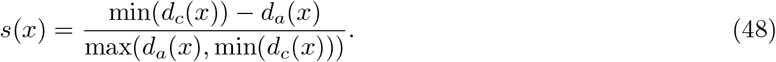

Then the batch silhouette score for the cell label *i* is defined as,

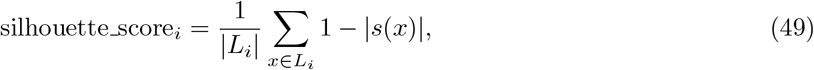

where *L*_*i*_ is the set of cells with the cell label *i* and |*L*_*i*_| denotes the number of cells in that set. Then the average silhouette score is defined as

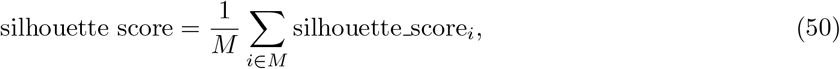

where *M* is the number of unique cell labels.

#### 8.3 HSIC on Batches

We utilize HSIC (see Section Hilbert Schmidt Independence Criterion (HSIC)) to evaluate how independent the latent representations are from batch variables. The lower the values are, the more independent the latent representations are from batch variables.

#### 8.4 Normalized Mutual Information (NMI)

Normalized Mutual Information (NMI) is a metric used to evaluate the similarity between two clustering assignments. It scales the Mutual Information (MI) score to a value between 0 and 1, where 1 indicates perfect agreement and 0 indicates no mutual information between the clusterings.

The NMI between two clusterings 𝒞 and 𝒞^′^ is defined as:

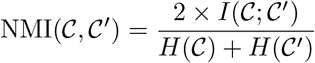

where:

- *I*(𝒞; 𝒞 ^*′*^) is the Mutual Information between 𝒞 and 𝒞 ^*′*^.
- *H*(𝒞) and *H*(𝒞 ^*′*^) are the entropies of 𝒞 and 𝒞^*′*^, respectively.

The Mutual Information *I*(𝒞; 𝒞 ′) is calculated as:

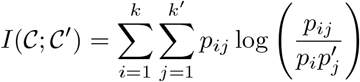

and the entropy *H*(𝒞) is:

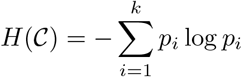

Similarly for *H*(𝒞^′^):

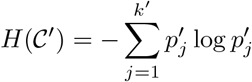

where:

- 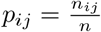 is the probability that a randomly chosen data point belongs to cluster *i* in 𝒞 and cluster *j* in 𝒞^′^.
- 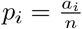 is the probability of cluster *i* in 𝒞.
- 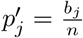 is the probability of cluster *j* in 𝒞^′^.
- *n*_*ij*_ is the number of data points in both cluster *i* and cluster *j*.
- *a*_*i*_ =Σ _*j*_ *n*_*ij*_ and *b*_*j*_ =Σ _*i*_ *n*_*ij*_ are the sums over rows and columns of the contingency table, respectively.
- *n* is the total number of data points.

#### 8.5 Definition of Adjusted Rand Index (ARI)

The Adjusted Rand Index (ARI) measures the similarity between two clusterings by considering all pairs of samples and counting pairs that are assigned in the same or different clusters in the predicted and true clusterings. It adjusts the Rand Index (RI) for the chance grouping of elements.

The ARI is defined as:

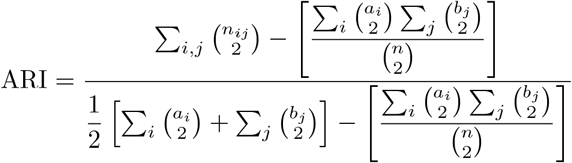

where:

- *n*_*ij*_ is the number of data points in both cluster *i* in 𝒞 and cluster *j* in 𝒞^′^.
- *a*_*i*_ = Σ_*j*_ *n*_*ij*_ is the number of data points in cluster *i* in 𝒞.
- *b*_*j*_ =Σ_*i*_ *n*_*ij*_ is the number of data points in cluster *j* in 𝒞^′^.
- *n* is the total number of data points.

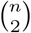 denotes the binomial coefficient, representing the number of ways to choose 2 items from *n*.

### 9 Optimal Transport for Lineage Prediction

#### 9.1 Optimal Transport

Given two probability vectors *r* and *c*, we consider the set of transport plans *U* (*r, c*):

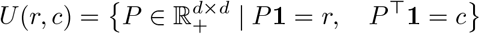

Here, **1** is a vector of ones. The set *U* (*r, c*) includes all non-negative *d* ×*d* matrices *P* where the sums of the rows equal *r* and the sums of the columns equal *c*.

The optimal transport distance between *r* and *c* using a cost matrix *M* and transport matrix *P* is defined as:

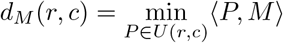

where ⟨*P, M*⟩ denotes the sum of the element-wise products of *P* and *M* . This represents the minimal total cost of transporting the distribution *r* to *c* under the cost *M* .

The Sinkhorn distance, introduced by [20], is a widely used method to make optimal transport computations more efficient by incorporating entropic regularization, which simplifies the optimization process. The dual problem of the entropic regularized optimal transport becomes:

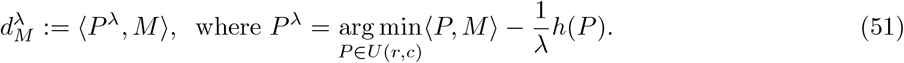

Here, λ *>* 0 is the regularization parameter, and *h*(*P*) denotes the entropy of *P* . Solving this regularized problem is computationally much cheaper than the classical optimal transport problem for suitable values of *>* [20]. In the following sections, we use SinkhornOT() to represent the function to compute *P* given *r, c*, and *M* .

#### 9.2 Lineage Prediction with Optimal Transport

We utilize optimal transport to further validate that the latent coupled/decoupled representations are able to predict lineage trajectory of cells. For each cell *i*, with *i* = 1, 2,. .., *n*, we have the (decoupled or coupled or both) latent representations **Z**_*i*_. Then we construct the cost matrix, where each element is the distance *d* between each cells. Formally, the cost matrix can be formulated as follows,

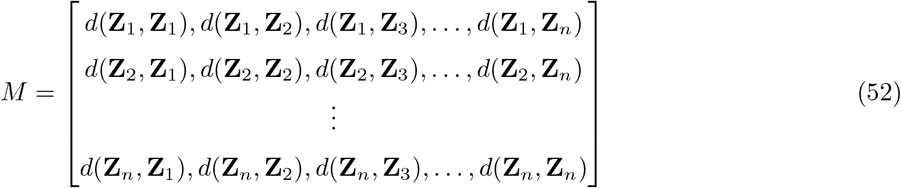

Then we solve the Optimal Transport problem using Sinkhorn Distance with the defined cost matrix *M*, initial cell states *r*, and the end cell states *c*. The goal is to compute the transportation matrix *P* . Specifically,

1. Compute *M* with Eq. 52.
2. Compute *c*, where 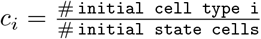 for all initial state cell types *i*.
3. Compute *r*, where 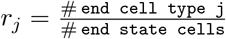 for all the end state cell types *j*.
4. Compute the transportation matrix *P* = `SinkhornOT`(*r, c, M*).
5. Aggregate the matrix *P* to get cell type transition matrix *T*, where 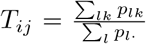, *l* is the cell index in cell type *i, k* is cell index in cell type *j*.

Finally we utilize the [POT](https://github.com/PythonOT/POT) v0.9.3 package to compute optimal transport.

### 10 Granger Causality-based Inference for Gene Regulatory Interactions

In this section, we introduce Granger causality and the Granger causal test, then we present the application of the Granger causal test to identify the distal, time-lagged regulation relations between the open chromatin regions and expression of nearby genes.

#### 10.2 Granger Causality and Tests

Granger [21] proposed a notion of causality based on how well past values of a time series *y*_*t*_ could predict future values of another series *x*_*t*_. Let *H*_*<t*_ be the history of all relevant information up to time *t −* 1 and *P* (*x*_*t*_|*H*_*<t*_) be the optimal prediction of *x*_*t*_ given *H*_*<t*_. Granger defined *y* to be causal for *x* if

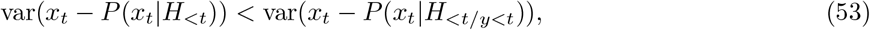

where *H*_*<t/y<t*_ is the optimal prediction that does not include the information of *y* up to time t. Granger causality assumes causal effects are ordered in time (i.e., cause before effect), under some assumptions: if *y* can predict *x*, then there must be a mechanistic (i.e., causal) effect; that is, predictability implies causality. Granger causality corresponds to nonzero entries in the autoregressive coefficients. In particular, for a bivariate model:

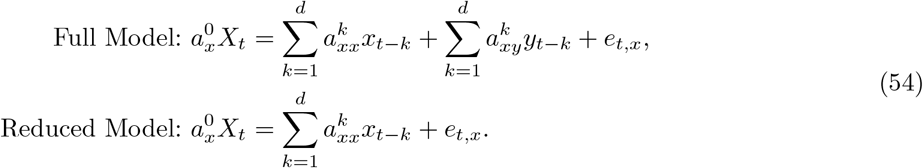

Series *y* is Granger causal for series *x* if and only if 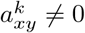 for some 1 ≤ *k ≤ d*. However, Granger causality based on only two variables severely limits the interpretation of the findings: Without adjusting for all relevant covariates, a key assumption of Granger causality is violated. In another words, Granger causality cannot resolve any latent confounders between two variables. Despite this limitation, Granger causal tests are still capable of revealing meaningful correlations between variables with time lagging information. Here we briefly introduce the Granger causal tests.

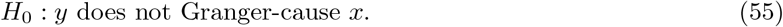

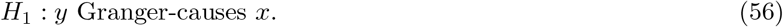

We can then construct the **F-test** test statistic

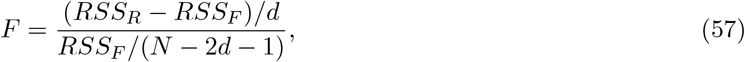

where *RSS*_*R*_, *RSS*_*F*_ are the residual sums of squares from the reduced and full models, respectively; *N* is the total number of samples; and *d* is the number of time lags [21]. If *F* exceeds the critical value from the F-distribution under the null hypothesis, reject *H*_0_. We can also utilize the **likelihood ratio test**,

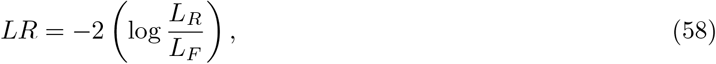

where *L*_*R*_, *L*_*F*_ are the likelihoods of the restricted and unrestricted models, respectively. Compute the *LR* statistic and compare it against a *p*-value from the chi-squared distribution with degrees of freedom equal to the number of time lags. If the *LR* statistic is greater than the *p*-value, reject *H*_0_.

#### 10.2 Granger Causal Regulatory Interaction Inference

In this section, we show how we utilize the Granger Causality test to reveal distal regulation with time lags. Specifically, we test whether peaks Granger-cause expression of nearby genes with time lags (Algorithm 1).

##### 10.2.1 Given Latent Time T

We introduce the algorithm to identify the Granger causal relations between a peak or some aggregated peak count 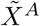 and gene expression of nearby gene 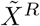 (for simplicity of notation, we drop the gene and cell indices here). In this test, we determine whether 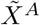 Granger-causes 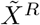.

###### Algorithm 1

Granger Causal Regulation Inference

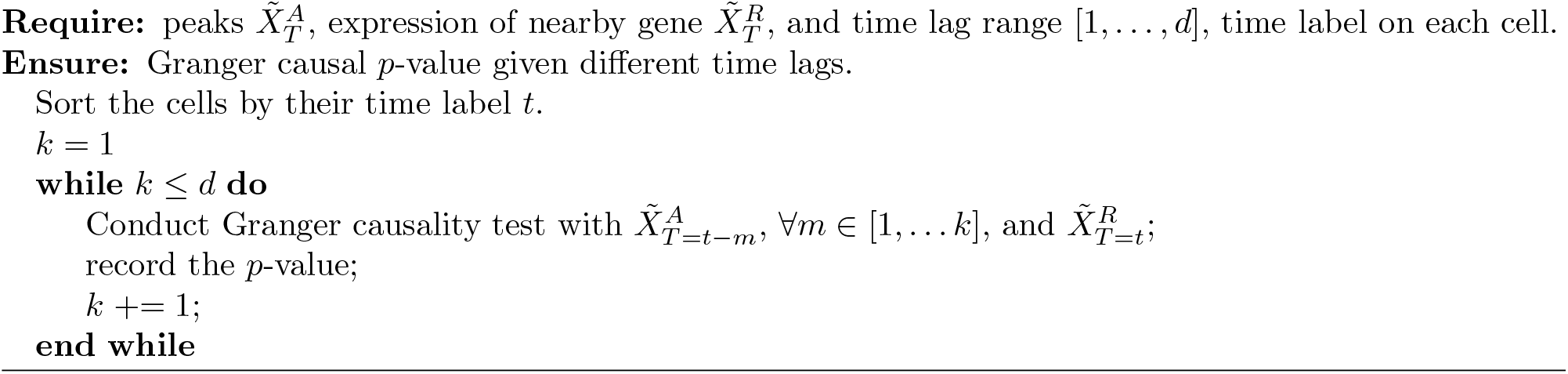

##### 102.2 Given Intra-nuclear Protein Information

The Granger causal regulatory interaction inference can be extended to use the protein levels of intra-nuclear TFs. By utilizing TF protein information as a proxy variable for latent time to order cells, the Granger causality test can be used to uncover TF-associated regulation by downstream enhancers of their gene targets.

### 11 Individual Gene Level Simulation

The simulation process consists of two main steps:

#### 1. Simulating scRNA-seq Gene Expression Data

For each gene *g* in cell *i*, we generate gene expression data using the Negative Binomial distribution:

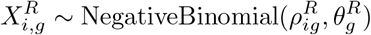

#### 2. Simulating Corresponding Peaks

For each peak *j* in cell *i*, we generate data using the Bernoulli distribution:

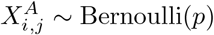

While a Multinomial distribution could also be used, we opt for the Bernoulli distribution here for simplicity, as suggested in [10].

Based on the previous sections, we establish the following causal relationships:

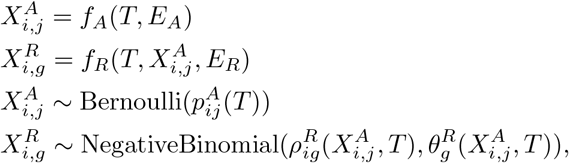

where *E*_*A*_ and *E*_*R*_ are the exogenous variables for gene expression and peaks respectively. The procedures of simulating time *T* and peaks 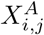 can be formulated as follows,

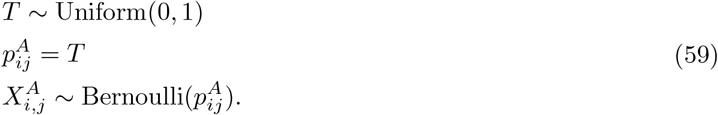

Note that here the parameter 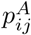 is a function of *T* (see section Problem Definition). Next, we simulate the gene expression 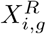 for the coupled and decoupled cases. For both coupled and decoupled cases,

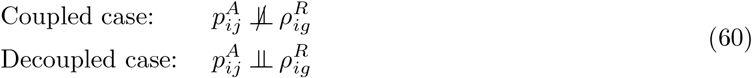

Specifically, for the coupled case, we simulate the following distributions,

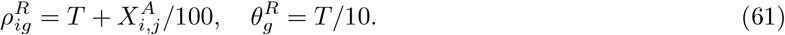

For the decoupled case, we simulate the following three distributions,

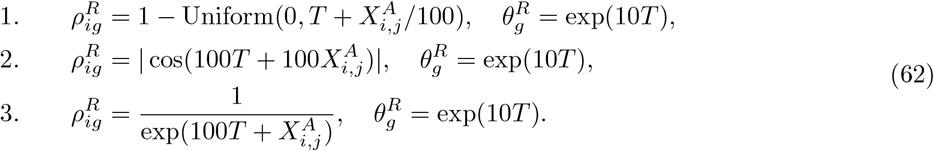

We repeat the simulation 100 times, with a sample size of 300 for each simulation.

### 12 Dataset Preprocessing and Settings

#### 12.1 SHARE-seq Mouse Skin (Hair Follicle) Dataset

We obtained this processed dataset directly from MultiVelo website[15]. There are total of 6,436 cells and 962 genes in the MultiVelo processed data. For the latent representations, the dimensions of latent coupled and decoupled representations are both 20. The Granger causality test was performed using likelihood ratio test to assess cis-regulation.

##### Algorithm 2

Gene-level Simulation

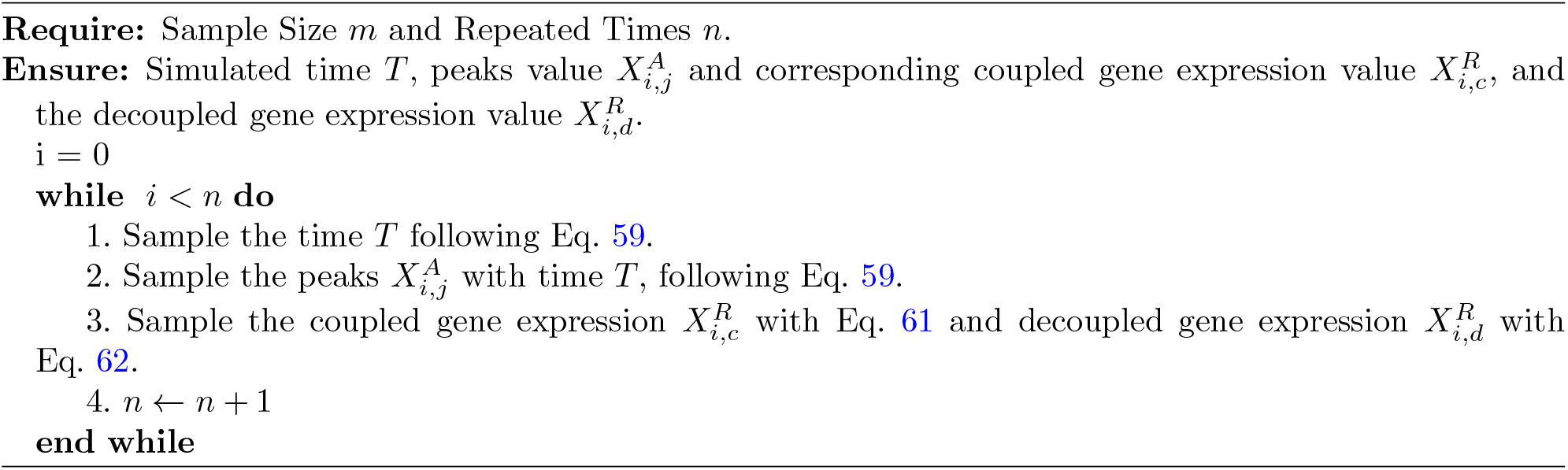

#### 12.2 NEAT-Seq Dataset

The NEAT-seq dataset profiles primary human CD4 memory T cells using a panel of master TFs that drive T cell subsets, including Tbet, GATA3, ROR,T, FOXP3, and Helios [16]. We obtained the processed dataset from GSE178707. After filtering out low-quality droplets, the dataset comprised 3370 cells with 13,380 genes and 78,203 peaks. The dimensions of the latent coupled and decoupled representations are both set to 30. The Granger causality test was performed using F-test explore gene regulation.

Binding sites of Tbet and GATA3 for Type 1 T helper (Th1) and type 2 T helper (Th2) cells were downloaded from Data S1 of https://doi.org/10.1038/ncomms2260 [22]. The set of Th1-specific T-bet super enhancers was downloaded from https://doi.org/10.1016/j.celrep.2016.05.054 [23].

To validate the effect of coexpressing T-bet and GATA3 on T cells compared to T cells expressing T-bet only, bulk RNA-seq data of EL4-Tbet+GATA3 and EL4-Tbet+Plum cells were downloaded from GSE171410.

#### 12.3 SSc Human Lung Epithelium Dataset

We used the 10X Genomics Multiome technology for paired snRNA-seq/snATAC-seq on nuclei from 6 SSc-ILD and 7 control human lung explants. For more details, please see [24].

4708 alveolar epithelial and terminal secretory cells, with 3000 highly variable genes and 178316 peaks, were used as input for the HALO model. Initially, the HALO model was trained based on ELBO loss only, and then the inferred latent representations were used for clustering and UMAP embedding construction using Scanpy [25]. Identified clusters were annotated using previously described cell markers [26]. Subsequently, the HALO model was trained using Palantir pseudotime for latent coupled and decoupled representations, where the dimensions of both are set to 20. The Granger causality test was performed using likelihood ratio test for distal regulation identification.

The spliced and unspliced RNA count matrices were generated using Cell Ranger output BAM files with Velocyto CLI (v0.17.17) [27]. These matrices were then processed with scVelo (v0.3.1) to compute RNA velocity using the “Dynamical” mode [28]. The RNA velocity-based cell-cell transition matrix was combined with cellular similarity to infer the initial and terminal states of cellular dynamics using CellRank (v2.0.2) [29]. Cells from the initial state were used as the root to compute pseudotime with Palantir (v1.3.2) [14].

To evaluate the information captured by HALO coupled and decoupled representations, a Multi-layer Perceptron classifier (scikit-learn v1.1.3) was trained to predict pulmonary epithelial cell type annotation.

Based on HALO representations, a cell-cell transition matrix was inferred using entropic-regularized optimal transport with a Sinkhorn solver (package POT v0.9.3) [20] and a cost matrix inferred based on squared Euclidean distance. This cell-cell transition matrix was then further aggregated into a cluster-cluster transition matrix. Additionally, we performed differential abundance testing to evaluate differences in cell abundances associated with disease using Milopy (v0.1.1) [30].

TF binding sites in the top important peaks were identified using FindMotifs function from the Signac package [31]. Association between top peaks and super enhancer sets was evaluated using the hypergeometric test, where super enhancer sets were downloaded from Data S1 of https://doi.org/10.1016/j.cell.2013.09.053 [32] and SEdb 2.0 [33]. We estimated DNA motif activity score using the Signac wrapper for chromVAR with motif profiles from the JASPER 2022 database [34, 35].

The scRNA-seq data of human lung organoids from GSE178360 were re-analyzed using Seurat and Harmony for normalization, batch correction, dimensionality reduction, and clustering [36, 37]. Marker genes and module scores for AT0, AT1, AT2, and TRB-SC were used to annotate cell types [26]. AT2 (Cluster 0) was isolated for differential gene expression analysis by comparing the EGF depletion group to the control group (Fig. S10). Differentially expressed genes (with absolute logFC *>* 0.5, min.pct *>* 0.2, and p.adj *<* 0.05) were used to calculate the EGF depletion module score for SSc lung epithelium data.

#### 12.4 NeurIPS Dataset

The NeurIPS single-cell multiomics dataset was collected from mobilized peripheral CD34+ hematopoietic stem and progenitor cells (HSPCs) isolated from four human donors at five time points. Samples were prepared using a standard protocol at four sites. The dataset was designed with a nested batch layout, with some donor samples measured at multiple sites and some donors measured at a single site. The processed data was downloaded from GSE194122. In our paper, we subset 10952 cells (Erythroblast, HSC, MK/E progenitor, Normoblast, and Proerythroblast) with pseudotime information for downstream analysis.The latent representation dimensions for scRNA-seq and scATAC-seq are both 10.

#### 12.5 Human Brain Dataset

We obtained this processed dataset directly from GSE162170. We followed MultiVelo’s protocol to process the dataset. The dimensions of the latent coupled and decoupled representations are both set to 20.

#### 12.6 10× Embryonic E18 Mouse Brain Dataset

We obtained this processed dataset directly from the MultiVelo website [15]. In total, it includes 3,365 cells, 936 highly variable genes, and 112,656 peaks; these matrices were then used for downstream analysis. The latent coupled and decoupled representation dimensions are both 10.

### 13 Enrichment Analysis

With the interpretable decoder of HALO, we can understand how a latent representation contributes to the reconstruction of gene or peak features. This allows us to obtain a list of gene or peak features contributing to a latent representation for enrichment analysis.

#### 13.1 Motifs

Using the JASPAR CORE collection, we compute motif profiles within scATAC-seq peaks using the MOODS algorithm [38], setting the adjusted p-value threshold to *p<* 1 × 10^*-*5^. Then, we identify TFs with predicted binding sites that are enriched in top peaks of ATAC latent representations **Z**^*A*^ (versus the remaining peaks) using the Fisher exact test.

#### 13.2 Genes

HALO uses Enrichr to find overlaps between the top genes from RNA latent representations **Z**^*R*^ and precompiled ontologies [39].

### 14 Gene Regulatory Network

We utilize the HALO representation to construct TF-CRE-gene linkages from scRNA/scATAC-seq data. For the regulatory network of AT2 cells, we use our larger independent scRNA-seq dataset (*n* = 17 SSc, 13 control lungs) to identify differentially expressed genes (DEGs) for the comparisons of interest [40].

#### 14.1 METAcell

Due to the sparsity inherent in single-cell genomics data, we aggregate individual cells exhibiting similar states into METAcells (see METAcell Construction for details).

#### 14.2 Identification of Cis-Regulatory Elements

To identify cis-regulatory elements (CREs), we adopt the nonlinear regression-based framework, DIRECT-NET [41]. Specifically, we use the non-linear predictive model XGBoost to regress the expression levels of genes against the accessibility values of candidate distal CREs, defined as peaks located within 250 kbp upstream or downstream of the transcription start site. CREs are selected based on the importance scores derived from the XGBoost model.

**Appendix Fig. 3:**
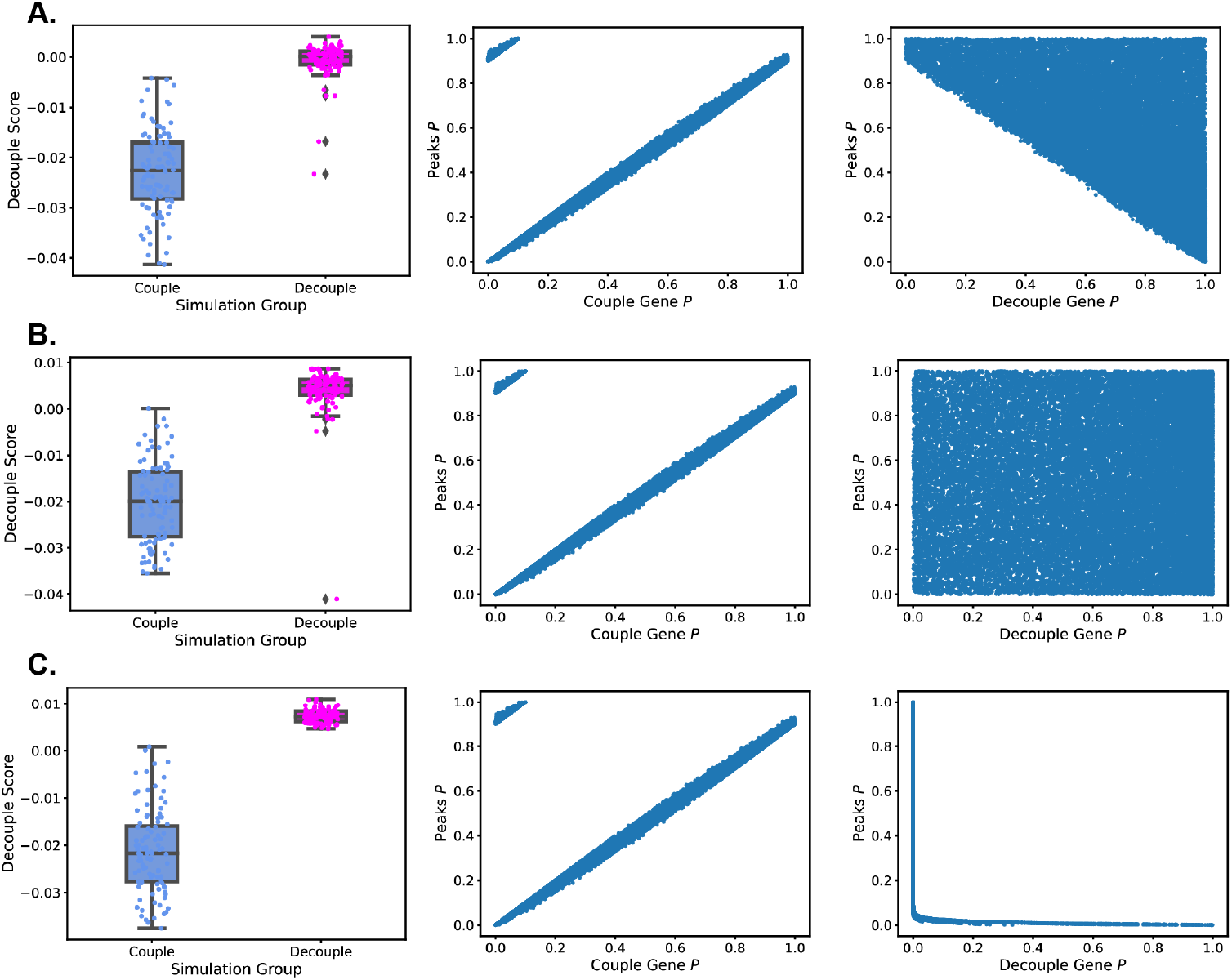
Simulation Results of Individual Gene Peak Pairs

#### 14.3 Construction of gene regulatory network

To construct a disease-associated regulatory network, we first identify DEGs (logFC *>* 0.5 and p.adj *<* 0.05) across conditions. Next, we identify differentially accessible CREs (logFC *>* 0.1 and p.adj *<* 0.05) corresponding to these DEGs. TFs within the differentially accessible CREs and promoter regions are identified using the JASPAR motif database and the motifmatchr function in the ChromVAR package [34, 35].

### 15 Training and Testing Setups

We extract 20% of the data from each dataset for testing. The remaining data is split into training and validation sets with a 4:1 ratio. For every dataset, we conducted an exhaustive hyperparameter grid search with n epoch=100 on the total loss with training and validation datasets. We denote dim as the dimension of the decoupled or coupled representations and we further set 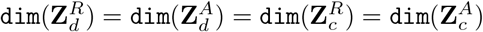. The search spaces for the hyperparameters [*ω*_1_, *ω*_2_, *α*, dim] are shown as follows:

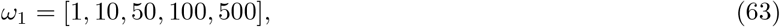

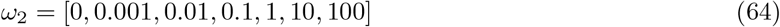

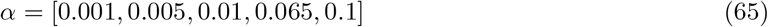

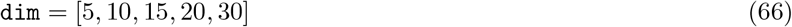

HALO is optimized using the Adam optimizer with a learning rate of 0.01, a weight decay of 0.001, and a minibatch size of 1024.

### 16 Ablation Study on Hyperparameters

#### 16.1 Hyperparameters Tuning

Here we examine the following hyperparameters: α in the coupled and decoupled constraints; *ω*_1_, the weight of causal constraints; and *!*_2_, the weight of the modality aligner. In the ablation studies for α, *ω*_1_, and *ω*_2_, we fix the latent representation dimensions as 10 for both scRNA-seq and scATAC-seq.

#### 16.2 α

The ablation study for different values of α was performed on two datasets, the mouse brain and human brain datasets, with α = [0.1, 0.065, 0.02] For this ablation study, we fixed *ω*_1_ = 10 and *ω*_2_ = 10. We report the values of 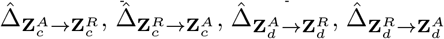, as well as 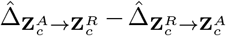 and 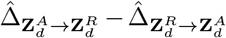 . We also report the clustering results for cell type annotation in terms of NMI and ARI in Appendix Fig. 4. From the results, we can see that generally, the results are most robust when α = 0.065.

#### 16.3 *ω*_1_

We conducted the ablation study of the causal constraints weights *ω*_1_ on the mouse brain dataset. *ω*_1_ are in the range of [1, 10, 50, 100, 500], while we fixed α = 0.065 and *ω*_2_ = 10. We report the values of 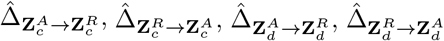, as well as 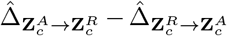 and 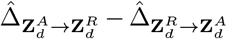 . We also report the clustering results for cell type annotation in terms of NMI and ARI in Appendix Fig. 5. From the results, we find that *ω*_1_ = 10 generally achieves the most stable performance.

#### 16.4 *ω*_2_

We conducted the ablation study of the modality aligner weights *ω*_2_ on the mouse brain dataset. *ω*_2_ are in the range of [0, 0.001, 0.01, 0.1, 1, 10, 100], while we fixed α = 0.065 and *ω*_1_ = 10. We report the values of 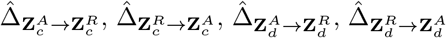, as well as 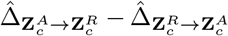 and 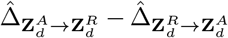 . We also report the clustering results for cell type annotation in terms of NMI and ARI in Appendix Fig. 6. From the results, we find that *ω*_2_ = 10 generally achieves the most stable performance.

#### 16.5 Number of Latent Representation Dimensions dim

We conducted the ablation study of the latent representation dimensions *n* on the mouse brain dataset. *N* are in the range of [6, 10, 18, 30], while we fixed α = 0.065, *ω*_1_ = 10, and *ω*_2_ = 10. We report the values of 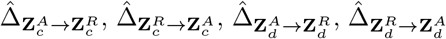, as well as 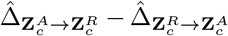 and 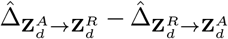. We also report the clustering results for cell type annotation in terms of NMI and ARI in Appendix Fig. 7. From the results, we find that *n* = 10 generally achieves the most stable performance.

**Appendix Fig. 4:**
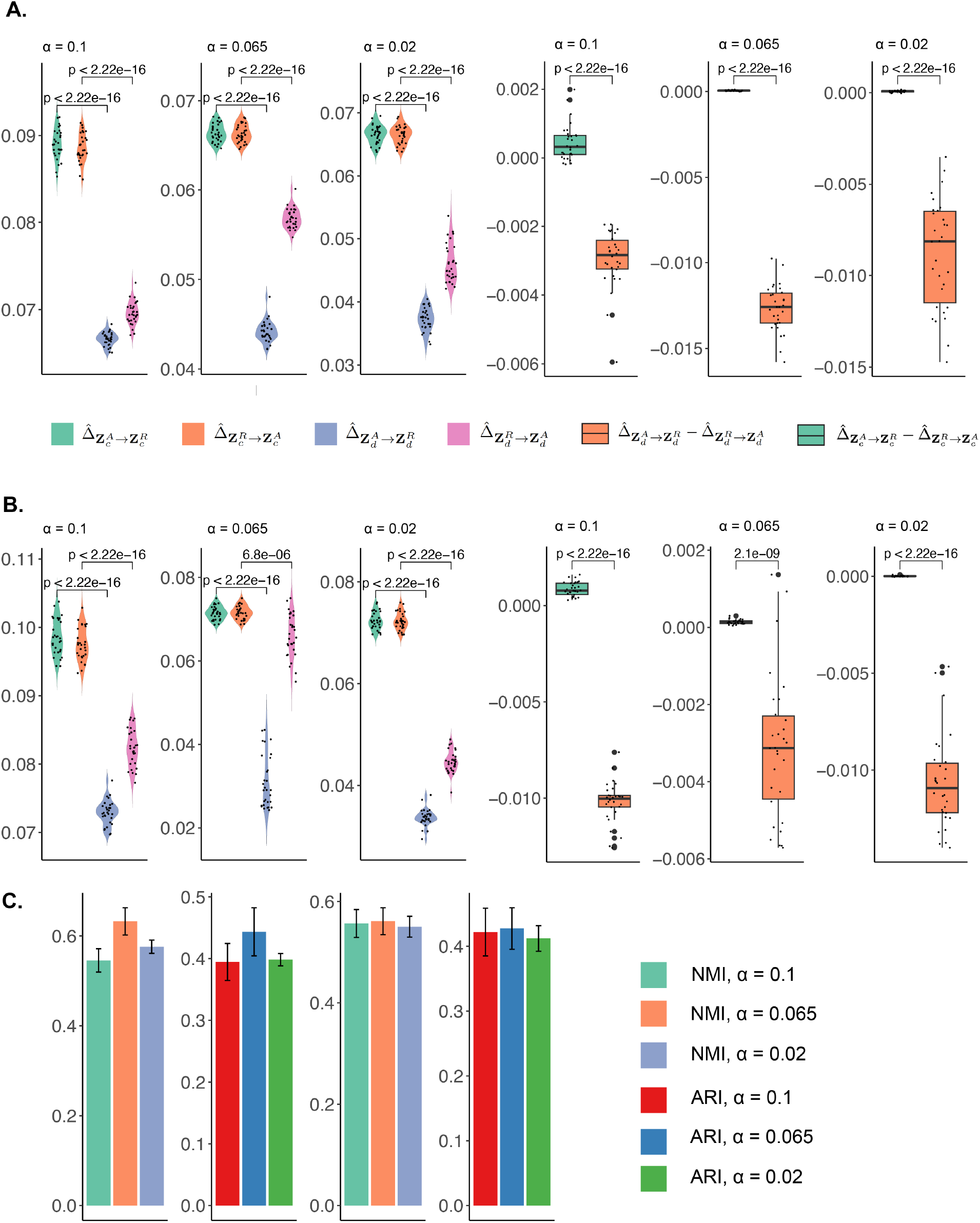
The ablation study with *α* = [0.1, 0.065, 0.02]. A. illustrates the results of mouse brain dataset. B. illustrates the results of human brain dataset. C. shows the clustering results of celltype annotation of mouse brain dataset in NMI and ARI.

**Appendix Fig. 5:**
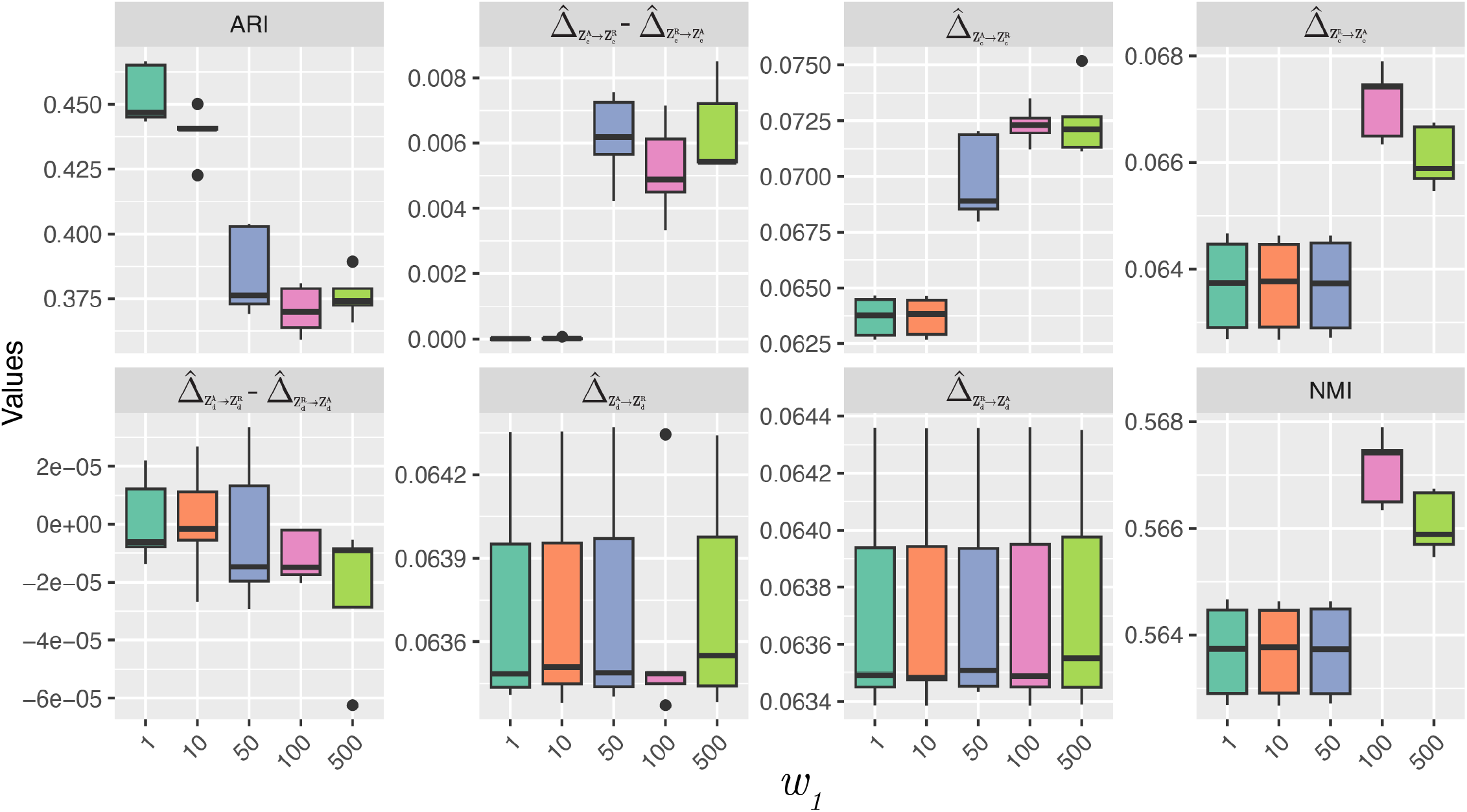
Ablation Study of *ω*_1_

**Appendix Fig. 6:**
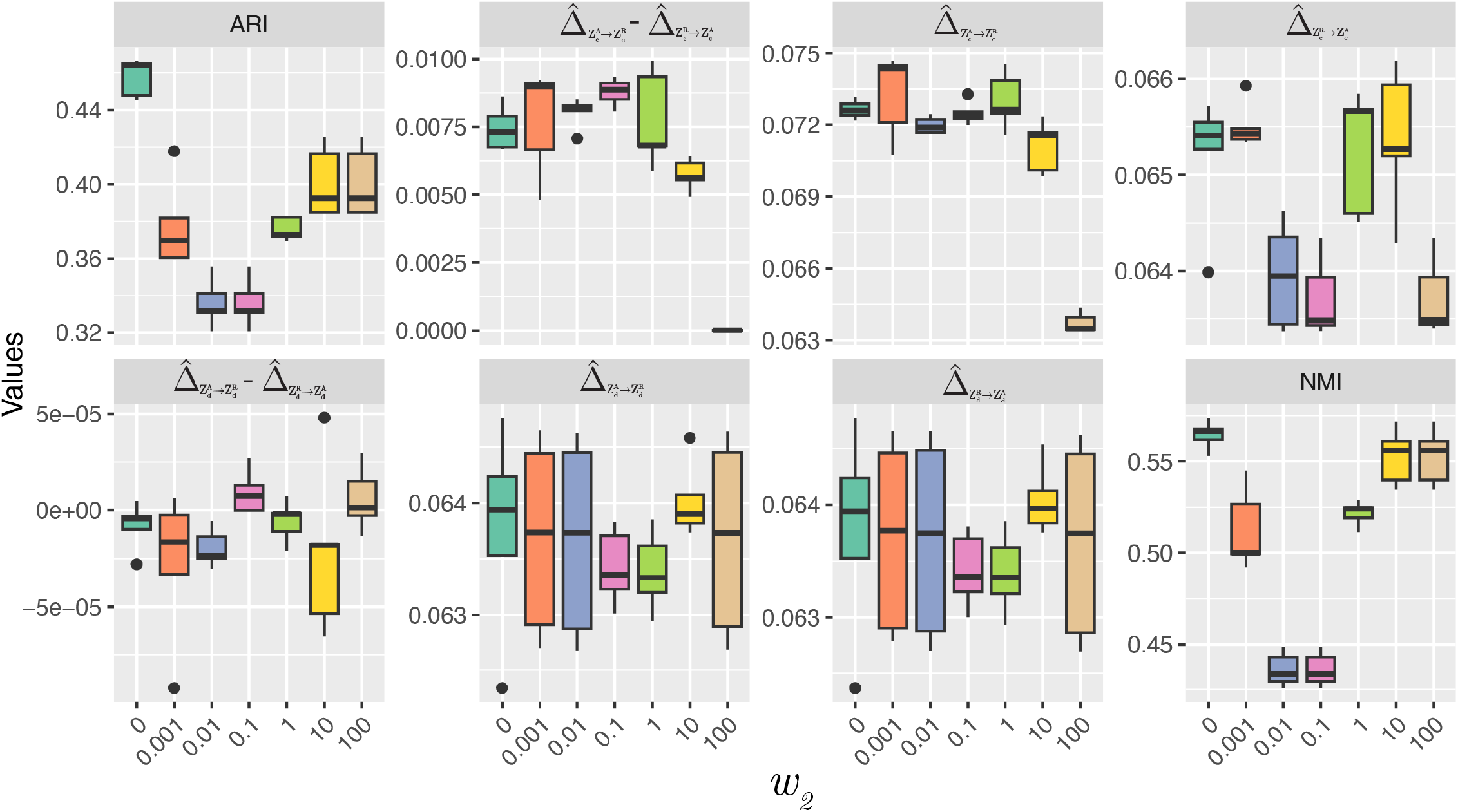
Ablation Study of *ω*_2_

**Appendix Fig. 7:**
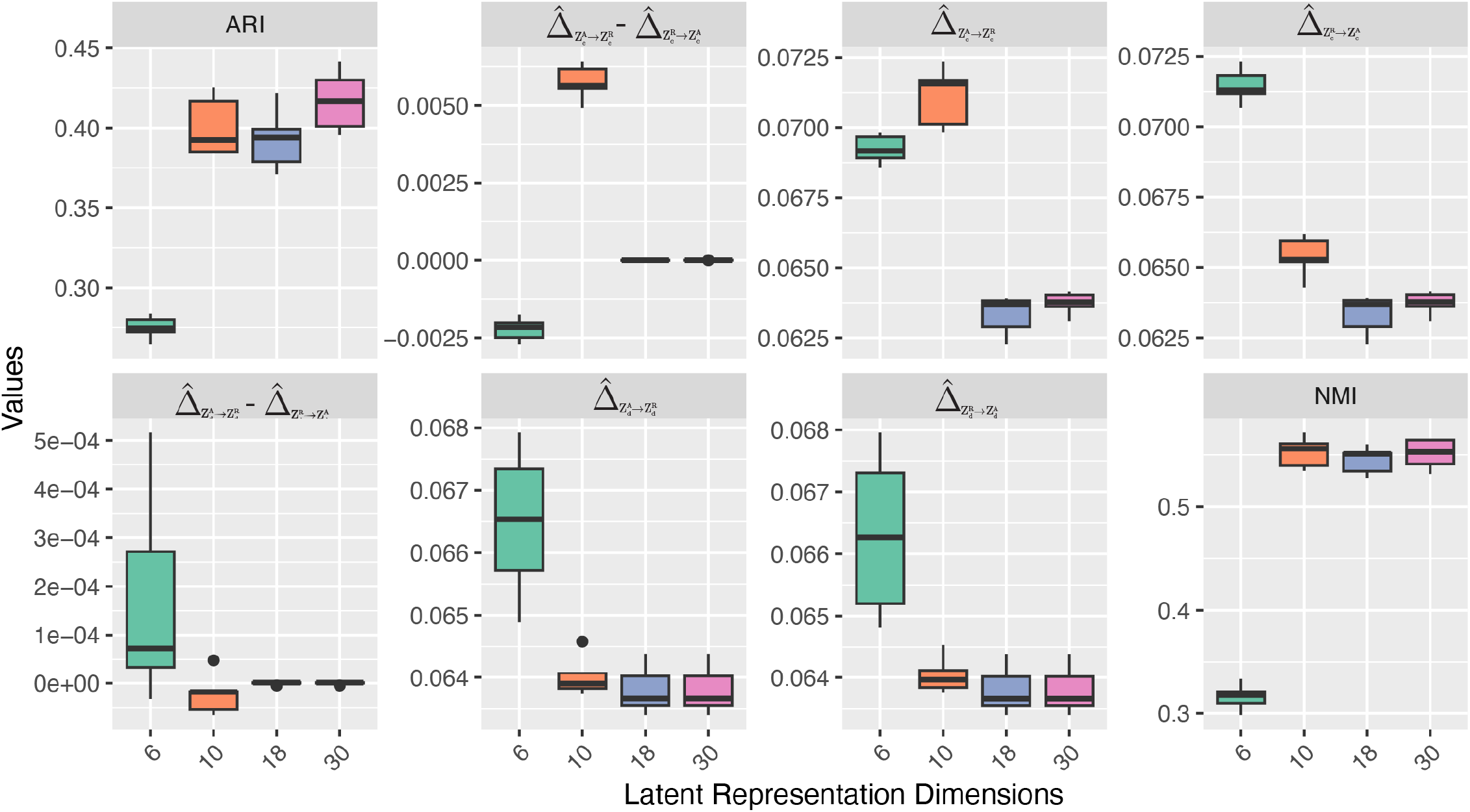
Ablation Study of latent representation dimensions

