## Extended Figure & table for "HALO: Hierarchical Causal Modeling for Single Cell Multi-Omics Data": Extended Figure.pdf

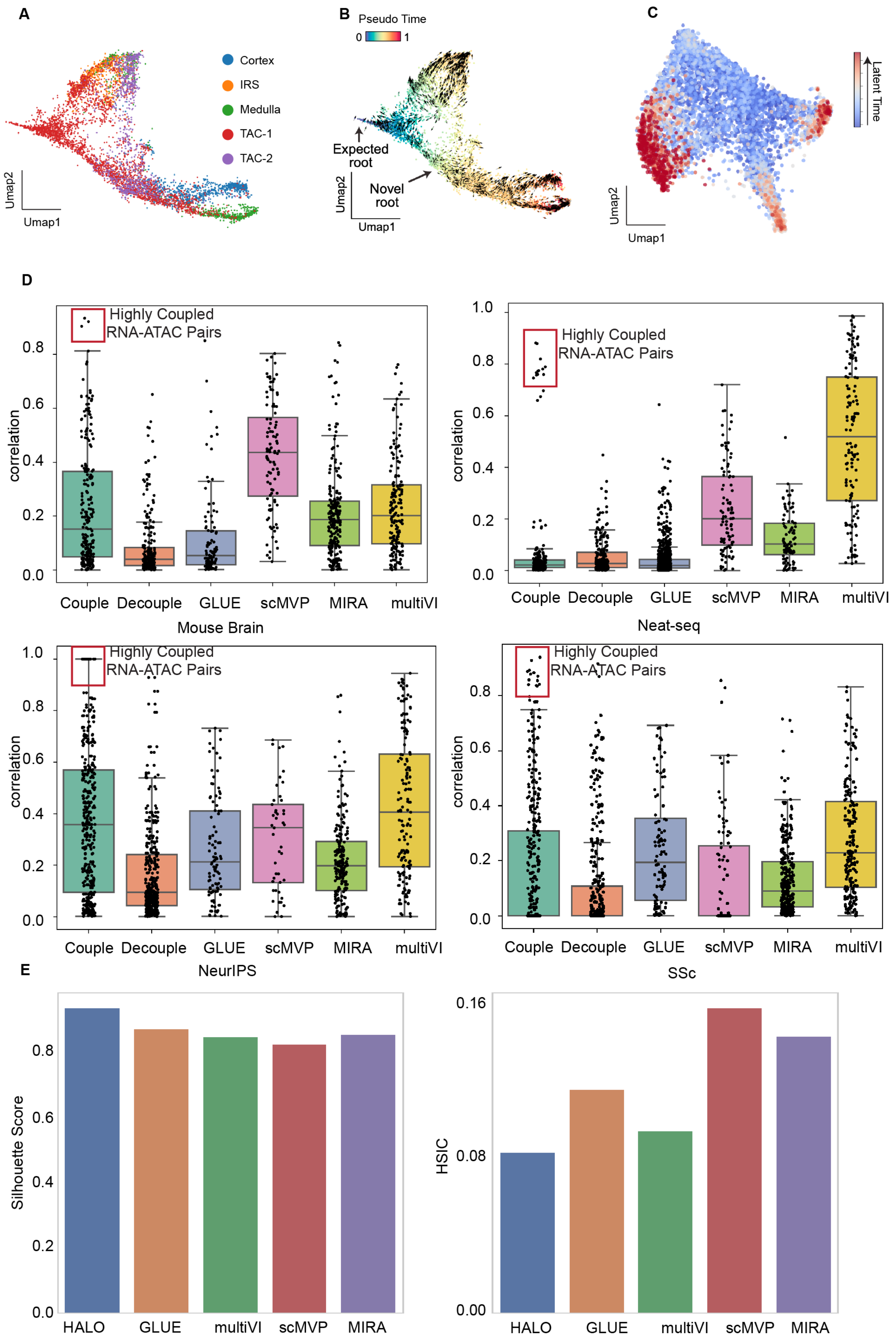
A
Cortex
IRS
Medulla
TAC-1
TAC-2
Umap2
Umap1
B
Pseudo Time
0
1
Expected root
Novel root
Umap2
Umap1
C
Latent Time
Umap2
Umap1
D
Highly Coupled RNA-ATAC Pairs
correlation
Couple
Decouple
GLUE
scMVP
MIRA
multiVI
Mouse Brain
Neat-seq
NeurIPS
SSc
E
Silhouette Score
HSIC
HALO
GLUE
multiVI
scMVP
MIRA

**Figure S1.** Mouse skin hair follicle data UMAP embedding and benchmark result.

A. The UMAP of original SHARE-seq mouse skin hair follicle dataset, colored by cell type.
B. The UMAP embedding of SHARE-seq data indicating novel root and expected root cells as identified by chromatin potential[8].
C. The UMAP generated from HALO concatenated RNA representations and ATAC representations, colored by the latent time estimated by MultiVelo[2].
D. The Pearson correlations among the representations of RNA and ATAC modalities for four datasets (mouse brain, NEAT-seq, NeurIPS, and SSc pulmonary epithelium).
E. The batch correction benchmarking on NeurIPS dataset. Left: the Silhouette Score (the higher the better), right: the HSIC of the representations and batch variables (the lower the better).

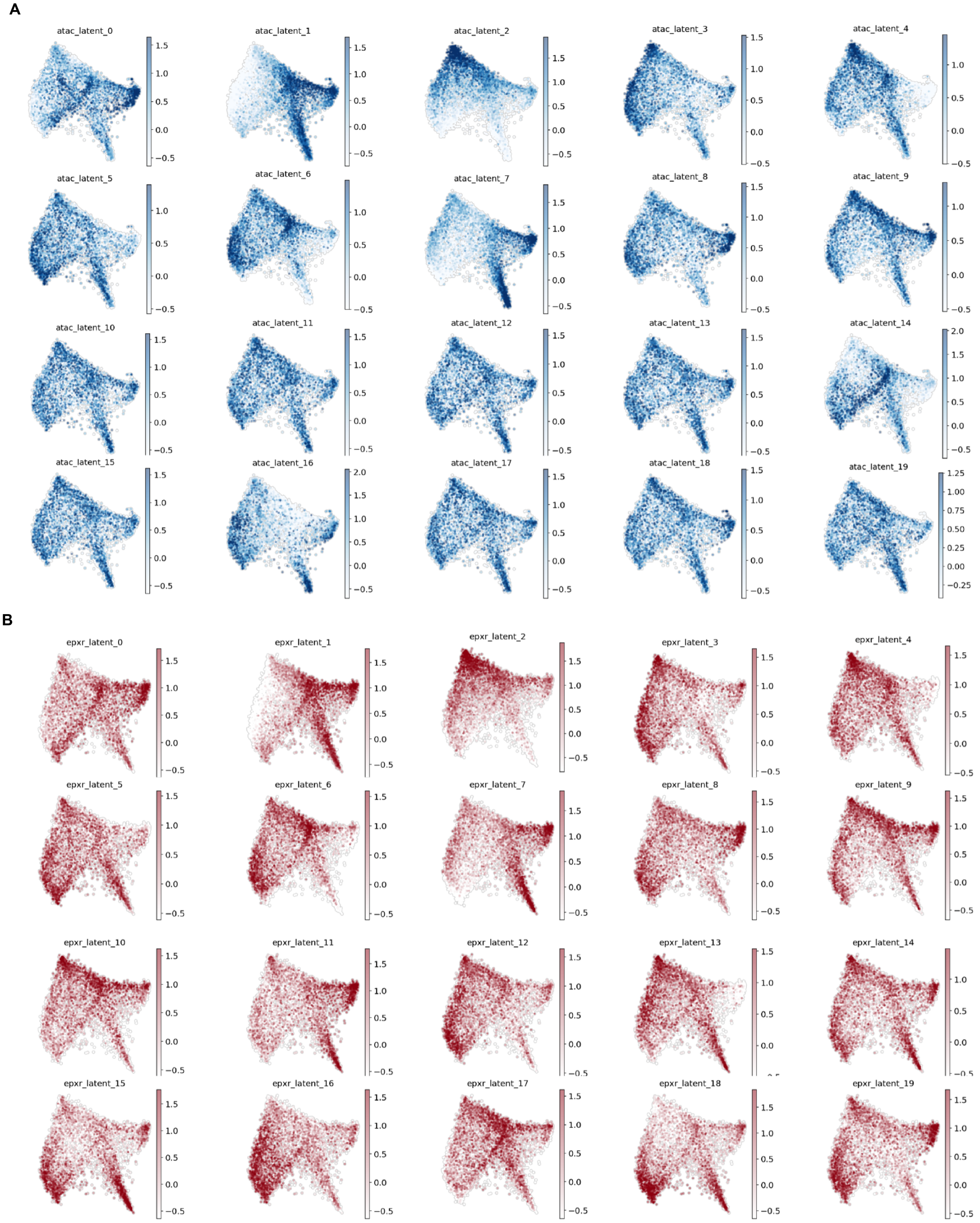
A
atac_latent_0
atac_latent_1
atac_latent_2
atac_latent_3
atac_latent_4
atac_latent_5
atac_latent_6
atac_latent_7
atac_latent_8
atac_latent_9
atac_latent_10
atac_latent_11
atac_latent_12
atac_latent_13
atac_latent_14
atac_latent_15
atac_latent_16
atac_latent_17
atac_latent_18
atac_latent_19
B
epxr_latent_0
epxr_latent_1
epxr_latent_2
epxr_latent_3
epxr_latent_4
epxr_latent_5
epxr_latent_6
epxr_latent_7
epxr_latent_8
epxr_latent_9
epxr_latent_10
epxr_latent_11
epxr_latent_12
epxr_latent_13
epxr_latent_14
epxr_latent_15
epxr_latent_16
epxr_latent_17
epxr_latent_18
epxr_latent_19

**Figure S2**. Distribution of the coupled and decoupled representations of RNA and ATAC data on UMAP embedding of the combined RNA and ATAC representations for the mouse skin hair follicle dataset.

A. ATAC latent representations. 0-10 are coupled ATAC representations, 11-20 are decoupled ATAC representations.
B. RNA latent representations. 0-10 are coupled RNA representations, 11-20 are RNA decoupled representations.

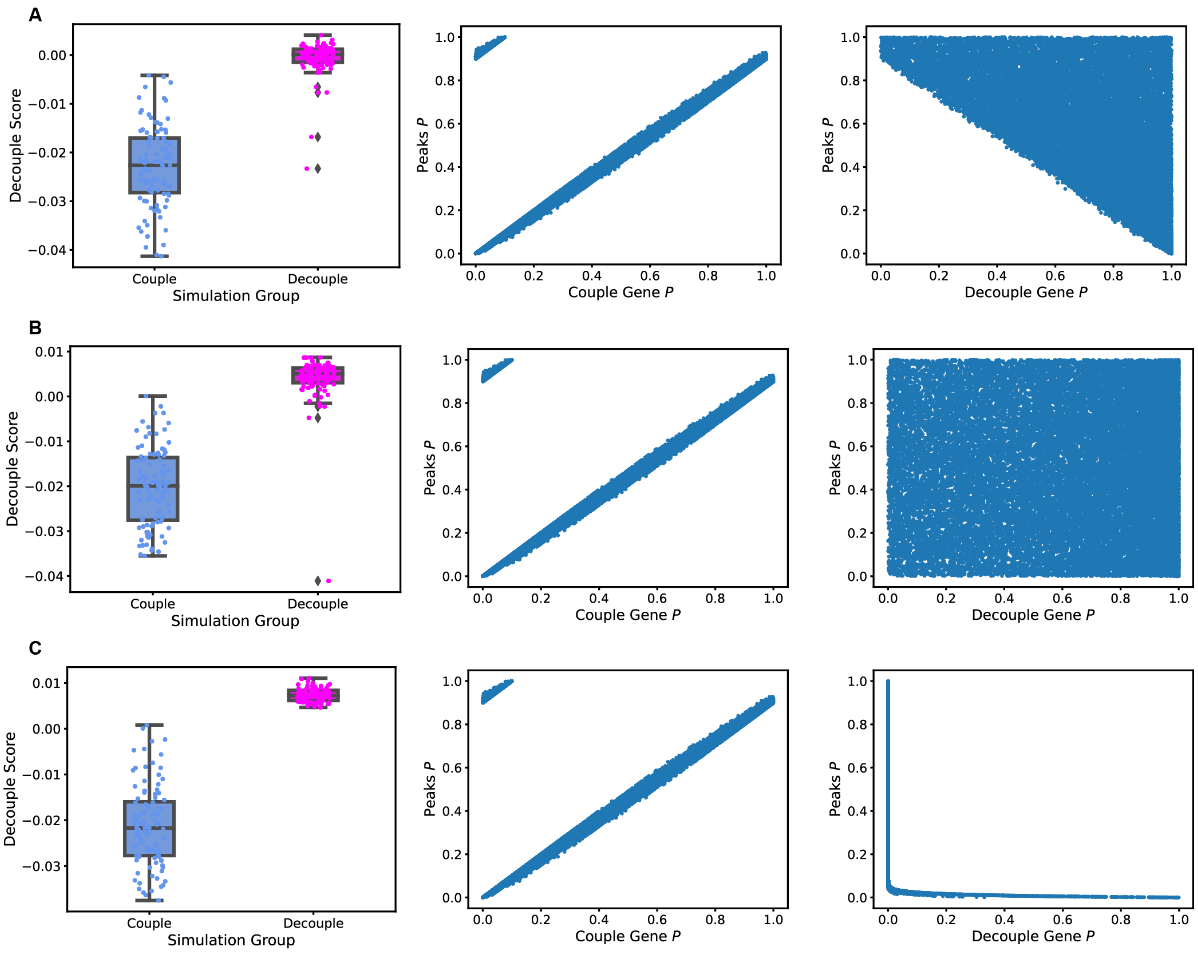


**Figure S3. Simulation results.** The simulation results of individual gene level (A) case 1, (B) case 2, and (C) case 3. The first column shows the decouple and couple simulation's decouple score; the second row shows relations between the coupled gene probability and corresponding peaks probability; the third row shows the relations of decoupled gene probability and corresponding peaks probability.

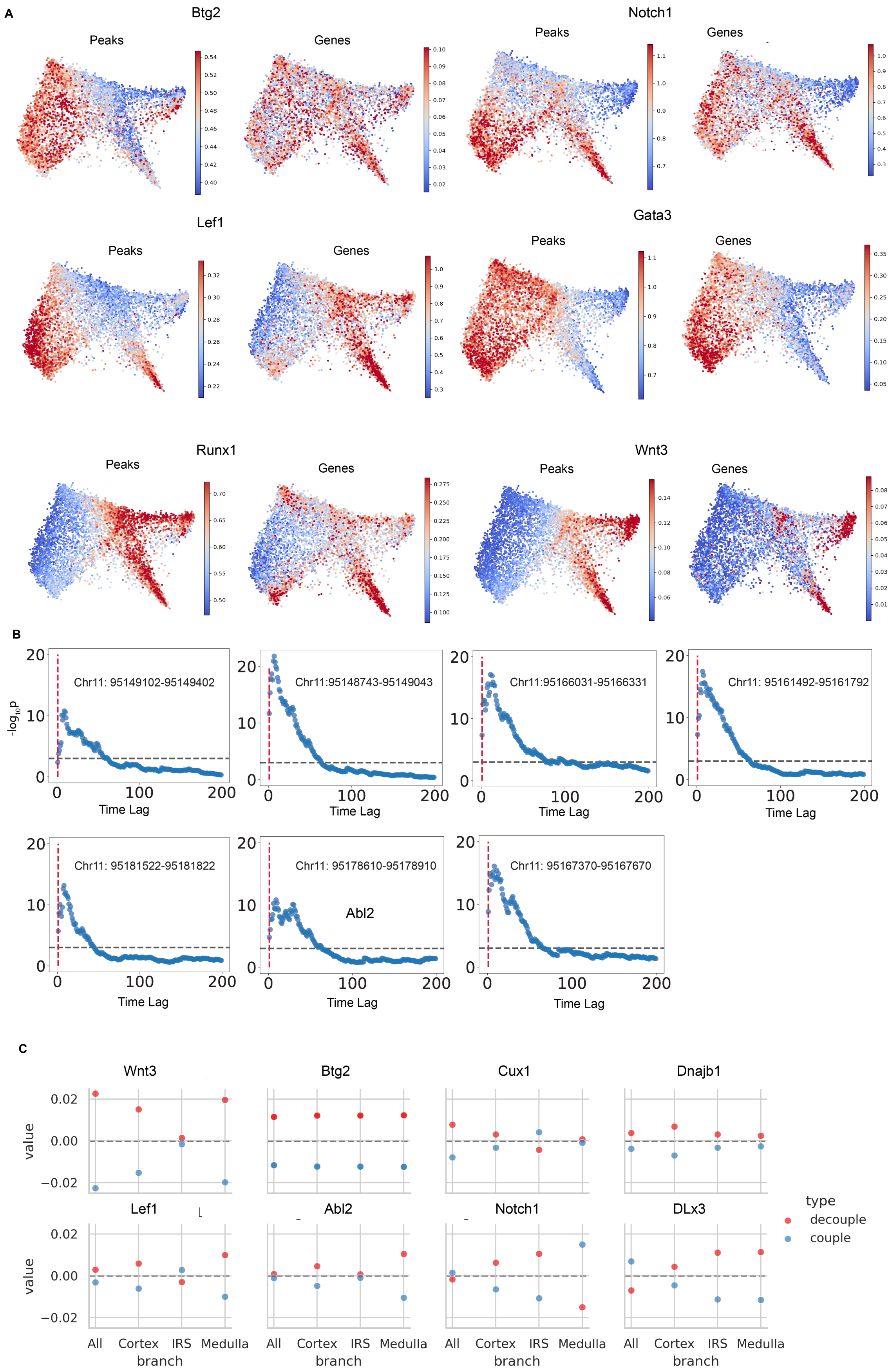

A
Btg2
Peaks
Genes
Notch1
Peaks
Genes
Lef1
Peaks
Genes
Gata3
Peaks
Genes
Runx1
Peaks
Genes
Wnt3
Peaks
Genes
B
-log10p
Chr11: 95149102-95149402
Chr11:95148743-95149043
Chr11:95166031-95166331
Chr11: 95161492-95161792
Chr11: 95181522-95181822
Chr11: 95178610-95178910
Abl2
Chr11: 95167370-95167670
Time Lag
C
Wnt3
Btg2
Cux1
Dnajb1
Lef1
Abl2
Notch1
DLx3
value
All
Cortex
IRS
Medulla
branch
type
decouple
couple

**Figure S4.** Gene-peak relation analysis in mouse skin hair follicle data.

A. Selected genes and their corresponding peaks.
B. The Granger causality test with local peaks of Dlx3 and gene expression of Itg2. The X-axis is time lag (number of cells, sorted by latent time). The Y-axis is the -log(P-value).
C. The couple and decouple score of selected genes on different branches.

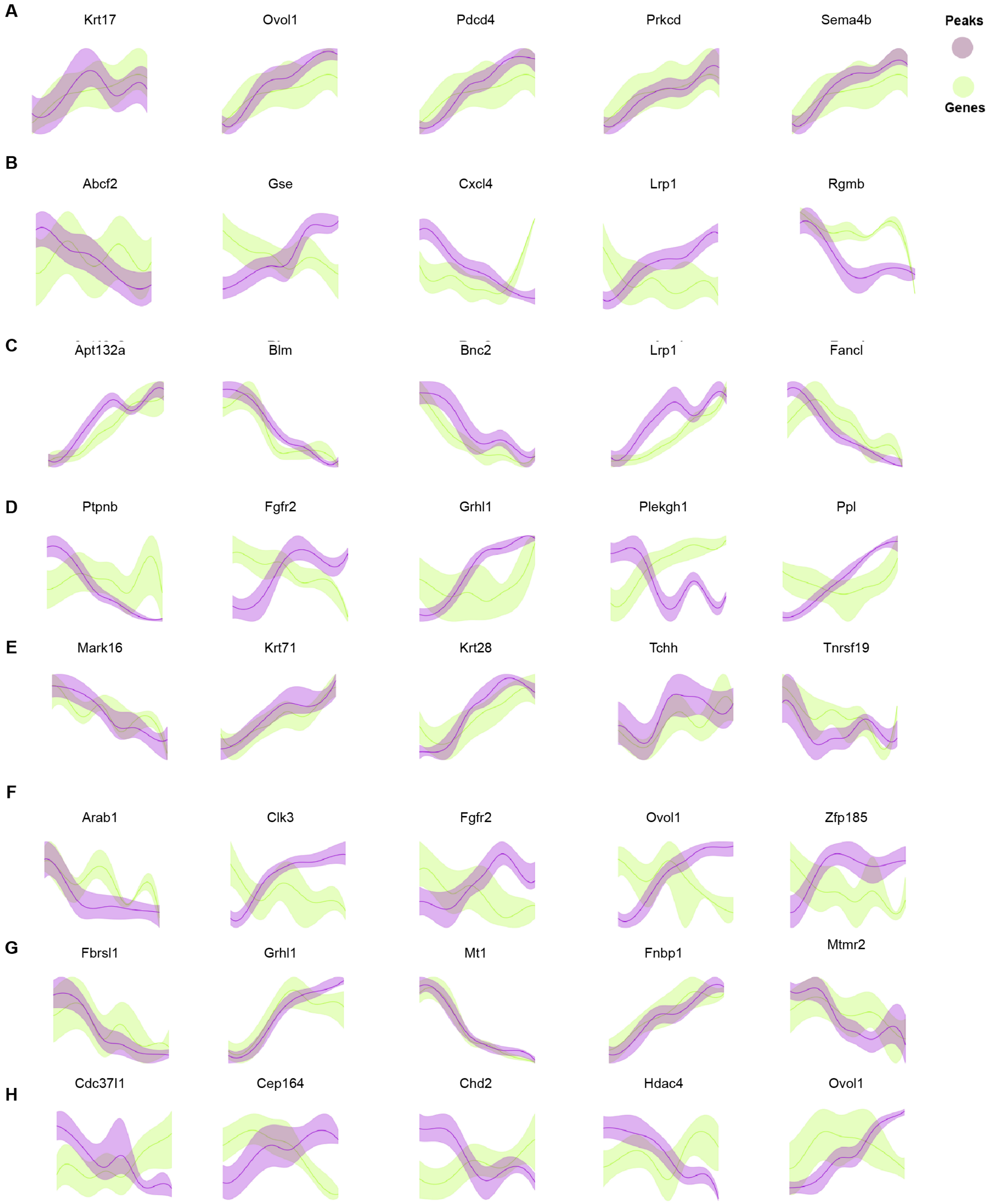


**Figure S5**. Coupled and decoupled genes on different branches.

A. Gene and peak levels that are coupled in all branches.
B. Gene and peak levels that are decoupled in all branches.
C. Gene and peak levels that are coupled in the Cortex branch.
D. Gene and peak levels that are decoupled in the Cortex branch.

E. Gene and peak levels that are coupled in the IRS branch.
F. Gene and peak levels that are decoupled in the IRS branch.
G. Gene and peak levels that are coupled in the Medulla branch.
H. Gene and peak levels that are decoupled in the Medulla branch.

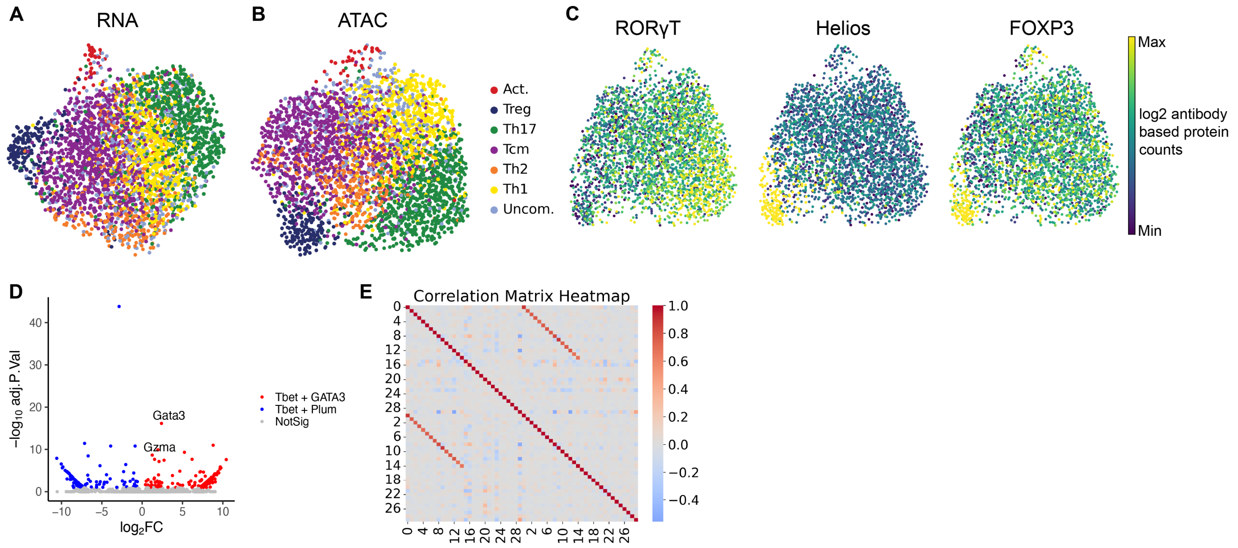


**Figure S6.** Additional figures for CD4+ effector T cell dataset.

A. UMAP embedding generated from HALO RNA representations.
B. UMAP embedding generated from HALO ATAC representations.
C. Nuclear protein levels of RORγT, Helios, and FOXP3 on UMAP of RNA and ATAC representations.
D. Volcano plot visualization of gene expression in EL4 cells expressing Tbet and GATA3 versus those expressing Tbet and Plum[23].
E. The Pearson-correlations matrix of latent representations.

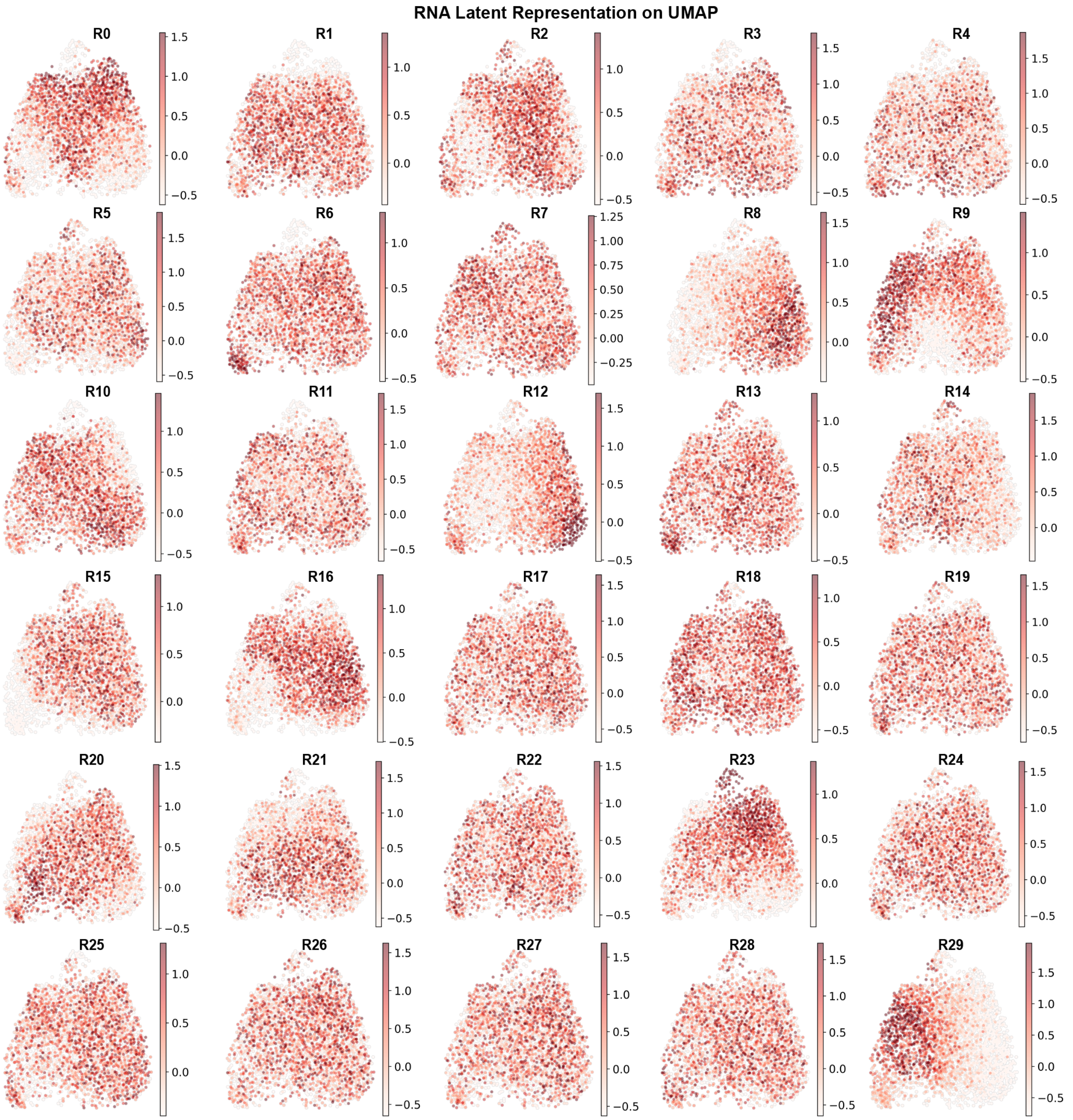


**Figure S7.** HALO RNA latent representation of human CD4+ effector T cells. 0-14 are coupled RNA representations, 15-29 are decoupled RNA representations.

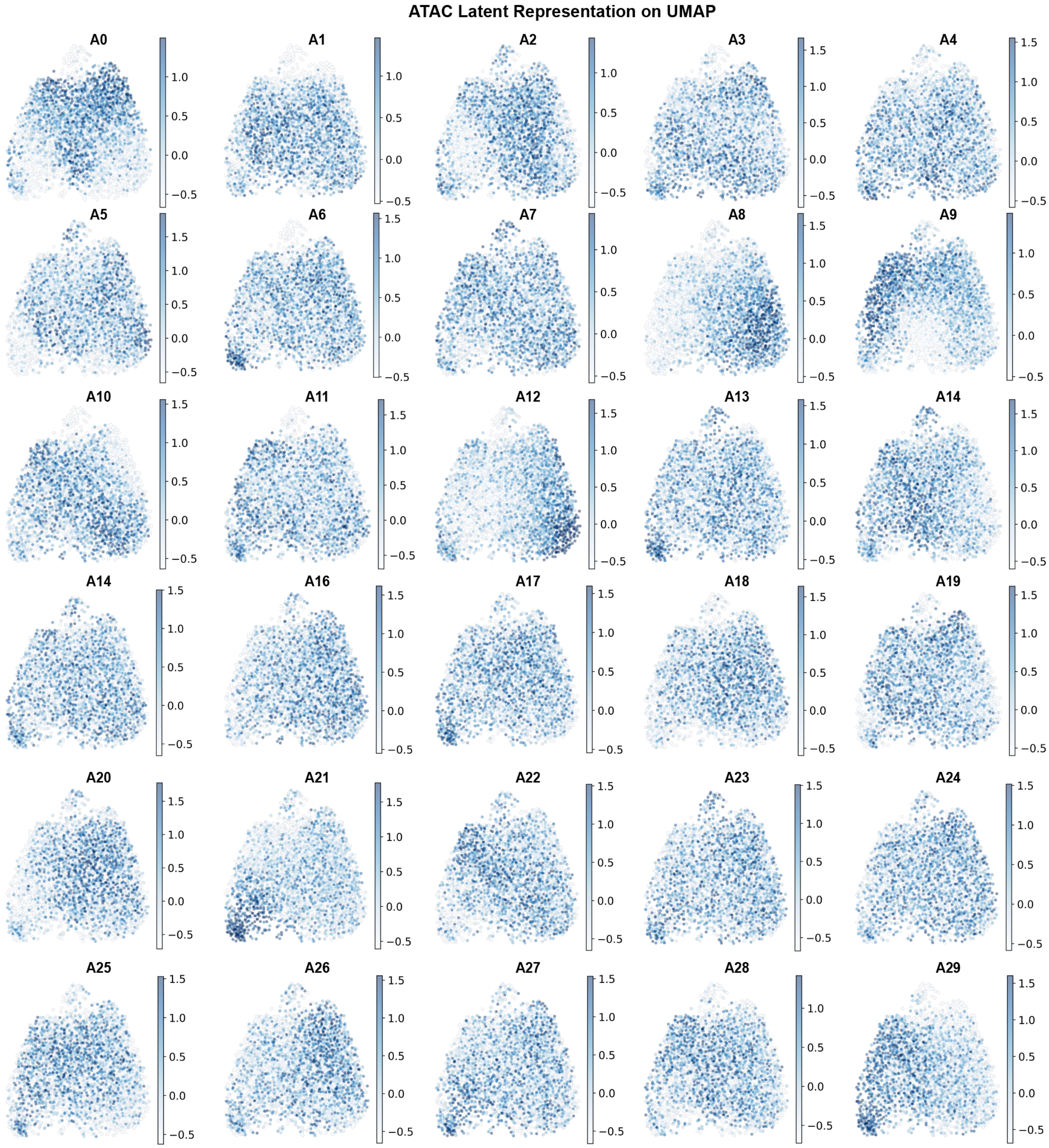


**Figure S8.** HALO ATAC latent representation of human CD4+ effector T cells. 0-14 are coupled ATAC representations, 15-29 are decoupled ATAC representations.

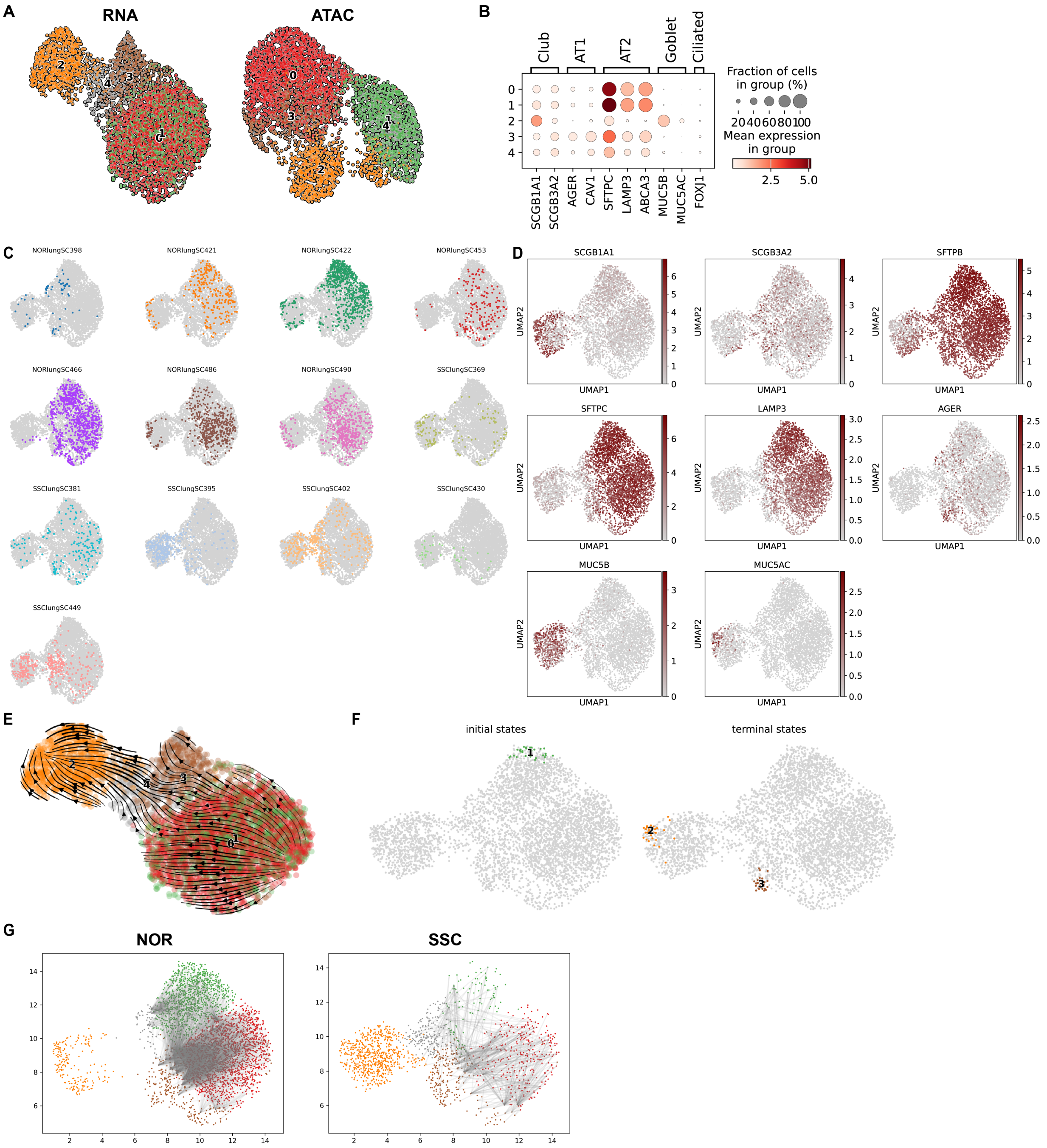


**Figure S9.** Additional figures for pulmonary epithelium dataset.

A. UMAP embeddings generated from RNA (left) or ATAC (right) representations.
B. Dot plot of marker genes for pulmonary epithelial populations.
C. Epithelial cells on UMAP embedding of RNA and ATAC representations, colored by sample.
D. Feature plots visualizing marker genes on UMAP embedding of RNA and ATAC representations.
E. Stream plot of RNA velocity on UMAP embedding of RNA representations.
F. Initial state and end states of cellular dynamics inferred by CellRank 2[30].

G. Optimal transport between initial state (AT2: clusters 0 & 1) and end states (clusters 2, 3, & 4). Top 0.1% edges with largest transition probability were visualized.

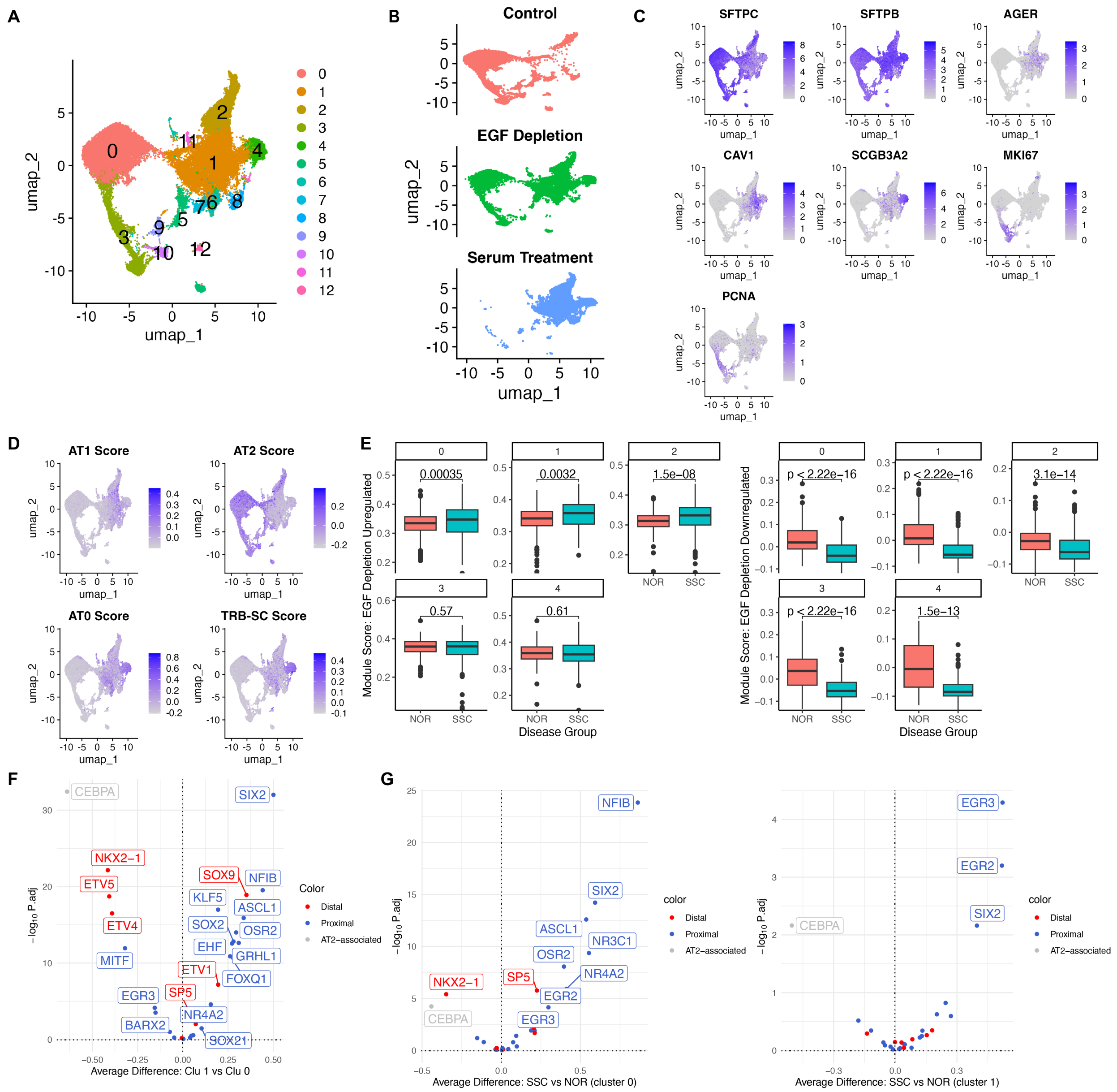


**Figure S10.** Additional results for pulmonary epithelium dataset.

A. Re-clustering of scRNA-seq data from human lung organoids under different treatments[27].

B. UMAP embedding of alveolar epithelial cells, colored by treatment.

C. Feature plots displaying marker gene expressions on the UMAP embedding.

D. Feature plots showing AT1, AT2, AT0, and TRB-SC scores on the UMAP embedding.

E. Module scores of EGF depletion in AT2 cells from human lung organoids. Scores were inferred based on differential gene expressions (DGEs) between EGF depleted and control cells in cluster 0 (AT2), as visualized in panel A.

F. Comparison of motif activity scores for transcription factors essential for proximal and distal airway patterning during development between AT2 subpopulations (clusters 0 & 1) from SSc pulmonary epithelium data.

G. SSc-associated changes in AT2 cells of motif activity scores for transcription factors essential for proximal and distal airway patterning during development.

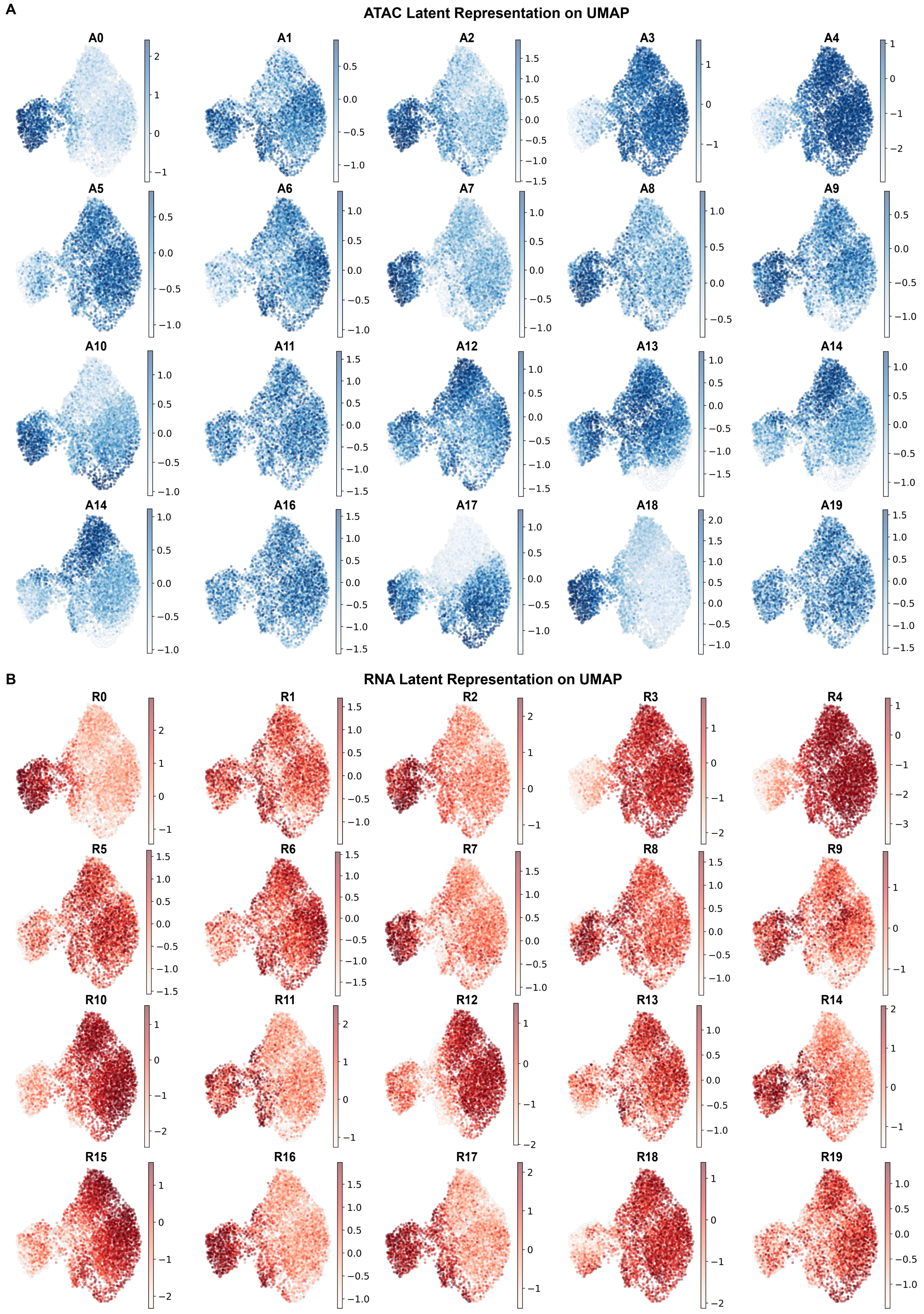
A
ATAC Latent Representation on UMAP
A0
A1
A2
A3
A4
A5
A6
A7
A8
A9
A10
A11
A12
A13
A14
A14
A16
A17
A18
A19
B
RNA Latent Representation on UMAP
R0
R1
R2
R3
R4
R5
R6
R7
R8
R9
R10
R11
R12
R13
R14
R15
R16
R17
R18
R19

**Figure S11.** HALO latent representations of pulmonary epithelial cells.

A. ATAC latent representations. 0-10 are coupled ATAC representations, 11-20 are decoupled ATAC representations.
B. RNA latent representations. 0-10 are coupled RNA representations, 11-20 are decoupled RNA representations.

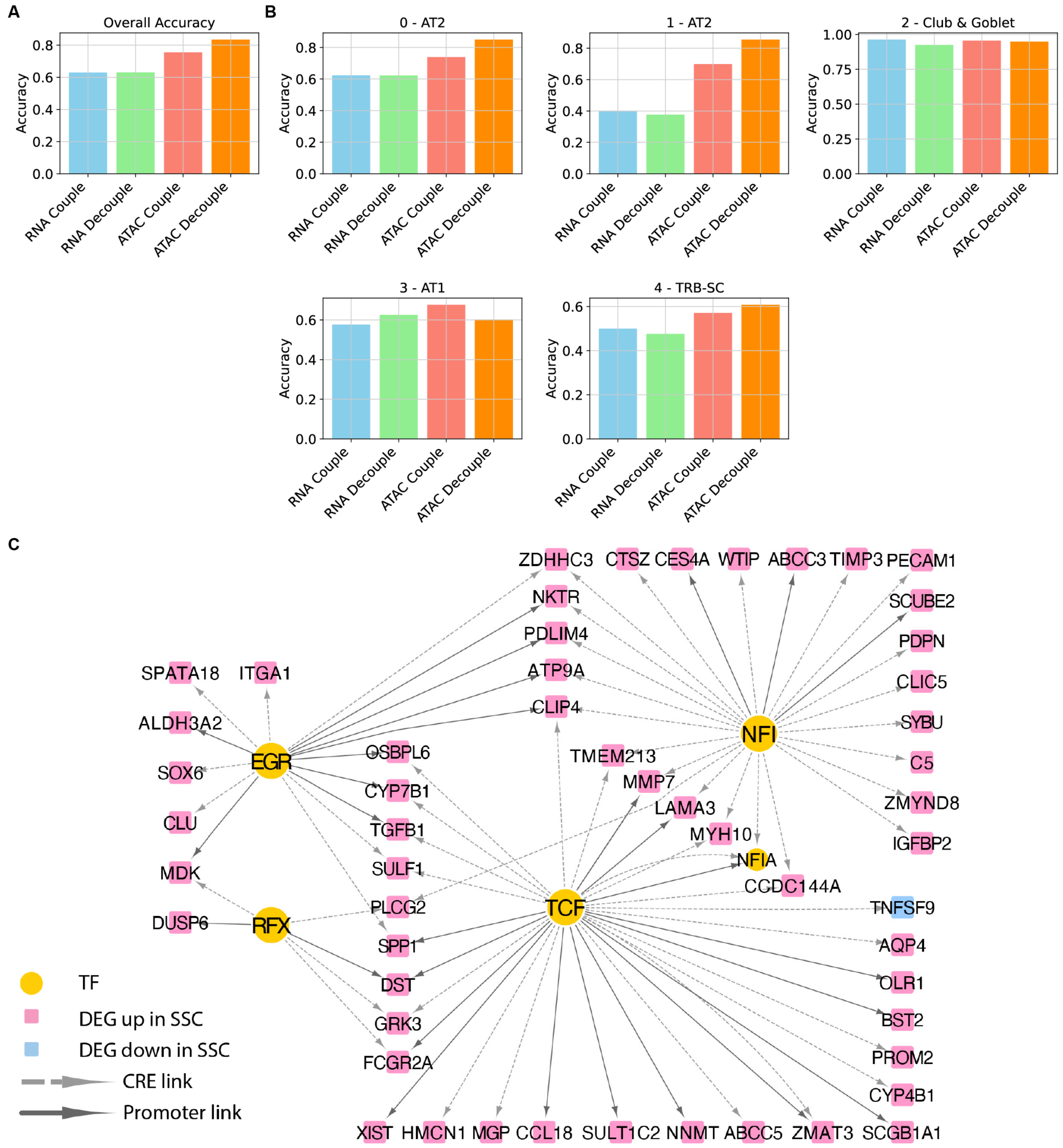


**Figure S12.** Pulmonary epithelial cell type prediction and SSc-associated gene regulatory network.

A. Overall accuracy of cell type prediction.
B. Cell type-specific accuracy for each cell type.
C. Gene regulatory networks for top enriched TFs (ranked by adj p-value) in SSc AT2 vs. Control AT2. Differentially expressed genes for this comparison were generated from a scRNA-seq dataset of 17 SSc and 13 control lungs[28]. Only DEGs with abslog2FC >1 are depicted.
